# Discovery of *Scrophularia nodosa* harpagoside synthase, a novel BAHD cinnamoyltransferase, bridges a key gap in the iridoid biosynthetic pathway

**DOI:** 10.64898/2026.04.10.711996

**Authors:** Dorian Rossi, Sheng Wang, Aude Pouclet, Yunyun Liu, David Pflieger, Etienne Grienenberger, Claire Parage, Ludivine Malherbe, Abdelmalek Alioua, Sandrine Koechler, Emmanuel Gaquerel, Danièle Werck-Reichhart, Nicolas Navrot

## Abstract

Harpagoside, a high-value anti-inflammatory iridoid compound, is traditionally extracted from the roots of *Harpagophytum procumbens* (Pedaliaceae, Lamiales), a Southern African desert plant widely used in traditional medicine but currently threatened by overexploitation. *Scrophularia nodosa* (Scrophulariaceae, Lamiales) is a perennial annual plant widely distributed in Western Europe and accumulates several iridoid compounds with known biological activities, such as catalpol, aucubin, harpagide and particularly harpagoside. We gathered extensive genomic and transcriptomics resources for this species and aimed at deciphering the biosynthetic pathway leading to the most abundant iridoid in *S. nodosa*, harpagoside. We found that the early iridoid pathway is well conserved with other iridoid-producing plants and validated the enzyme activities by transient co-expression in *N. benthamiana*. Investigation into the large BAHD family showed subclade 6i expanding in Scrophulariaceae, with an atypical VYPWG motif instead of the canonical DFGWG. In this branch, we discovered and characterized harpagoside synthase, a BAHD-type cinnamoyl transferase enzyme showing unique high specificity to the uncommon cinnamoyl-CoA acyl donor and catalyzing the final step of harpagoside biosynthesis. These results establish *S. nodosa* as a new model to investigate unexplored branches of the iridoids metabolism, and are a first step towards sustainable harpagoside and high-value cinnamoyl-containing conjugates production.

## Introduction

As precursors or bioactive molecules themselves (Ghisalberti 1998), the monoterpene-derived bicyclic compounds iridoids have attracted considerable research interest. Several reports have shown their valuable bioactivities, both in complex plant extracts used in traditional medicine and isolated molecules (Davini et al. 1986; Hamburger et al. 1991; Jin et al. 2008). Among them harpagoside is described as an important contributor to the anti-inflammatory and analgesic activity of the roots of a desert plant extensively used against osteoarthritis, *Harpagophytum procumbens,* as supported by its *in vitro* inhibition of the expression of cyclooxygenase-2 and inducible nitric oxide (Huang et al. 2006).

The biosynthesis of iridoids is well understood with regard to the “early” steps forming conserved structural features in this group of molecules, and notably the intermediate nepetalactol. This specialized pathway is initiated with the formation of the monoterpenol geraniol by a geraniol synthase. Geraniol then undergoes oxidations by the cytochrome P450 geraniol-8-hydroxylase (G8O) (Collu et al. 2001) and the oxidoreductase 8-hydroxygeraniol oxidase (8HGO) (Geu-Flores et al. 2012). The resulting product is cyclized by the iridoid synthase (ISY) to form nepetalactol (Munkert et al. 2015). Issued from this common intermediate, a limited number of iridoid “cores” have been described in plants, and the huge diversity of iridoids found in nature, surveyed in extensive reviews (Dinda et al. 2007a, 2007b, 2009, 2011) mainly originates from “decorations” of these cores, including extensive glycosylation, acylation, and/or oxidation at various positions. Core variants differ in the structure of the cyclopentanopyrane (iridane) ring. In the most common molecules, the presence of either an epoxide or a double-bond in the pentane ring between carbons 7 and 8 and leads to catalpol and aucubin derivatives, respectively, whereas an unmodified single bond between carbons 7 and 8 give rise to harpagide-related molecules, found notably in *Scrophularia nodosa* (Figure 1, A, Figure 1, B). In some species, enzyme-catalyzed opening of the pentane ring leads to secoiridoids, commonly represented by secologanin, the precursor of complex indole alkaloids such as vinblastine found in *Catharanthus roseus* (Miettinen et al. 2014; Kulagina et al. 2022), or oleuropein, accumulated in olive *Olea europaea* L. (olive) fruit (Calderini et al. 2026) (Figure 1, A). Shorter biosynthetic pathways can lead to the production of small volatile bioactive iridoids such as nepetalactone, the pathway of which has recently been fully elucidated in *Nepeta cataria* (catnip) (Lichman et al. 2020) and *Teucrium marum* (cat thyme) (Smit et al. 2024).

**Figure 1.**
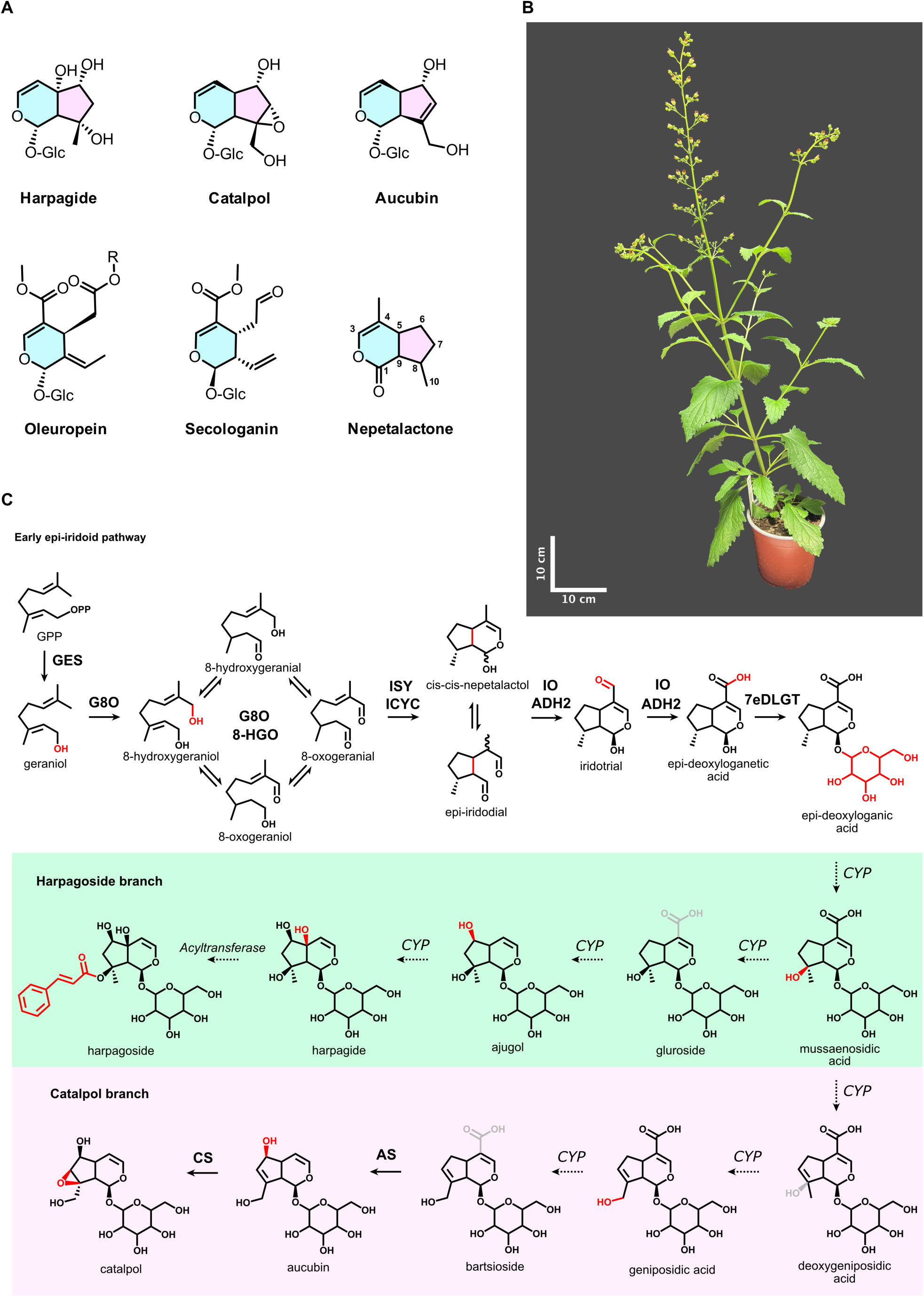
Iridoid pathway in *S. nodosa*. **A.** Structures of representative iridoids found in plants with conventional carbon numbering of the iridoid core displayed on the nepetalactone molecule (in blue: pyrane ring, in purple: pentane ring). Glc = glucose, R = C_8_H_10_O_2_. **B.** Representative specimen of *Scrophularia nodosa* at flowering stage in controlled conditions. **C.** Harpagoside and catalpol biosynthetic pathways. Note that both compounds derive from *cis-cis*-nepetalactol (and thus *epi*-iridotrial) isomer. Italics: putative enzyme family involved: cytochrome P450 (*CYP*) and acyltransferase. GPP: geraniol diphosphate, GES: geraniol synthase, G8O: geraniol-8-oxidase, 8-HGO: 8-hydroxygeraniol oxidase, IO: iridoid oxidase, ADH2: alcohol dehydrogenase 2, 7eDLGT: 7-deoxy-*epi*-loganetic acid glucosyl transferase, CS: catalpol synthase, AS: aucubin synthase.

Beyond the minimal set of enzymes required to produce detectable amounts of products, more proteins, such as alcohol dehydrogenases and cyclases (Brown et al. 2015; Colinas et al. 2025), have been shown to largely enhance the enzymatic efficiency of specific steps and are likely involved in the optimal and regulated functioning of the pathway *in vivo*. For instance in the early part of the iridoid pathway, iridoid cyclase (ICYC) was recently discovered and shown to control the stereospecific cyclisation of the enol intermediate to form nepetalactol intermediates (Colinas et al. 2025), the stereochemistry of which defines two mutually exclusive class of molecules in plants, iridoids and epi-iridoids. Specialized “late” decoration cytochromes P450 aucubin synthase (AS) and catalpol synthase (CS) catalyzing aucubin and catalpol final biosynthetic steps have also recently been characterized in *Rehmannia chingii* and in different Lamiaceae species, respectively (Figure 1, C) (Rodríguez-López et al. 2022; Wang et al. 2025).

In most iridoid-producing species, the nepetalactol intermediate is oxidized to deoxyloganetic acid (or its *epi*-isomer) by the cytochrome P450 iridoid oxidase (IO) and an alcohol dehydrogenase 2 (ADH2) before being glucosylated by 7-deoxy-*epi*-loganetic acid glucosyl transferase (e7DLGT), presumably enhancing the stability of the intermediate. It may then be subjected to one or more additional hydroxylations, putatively catalyzed by cytochrome P450 enzymes, which create anchor points for further decorations such as addition of one or more sugars, acyl chains and/or phenolic groups, depending on the final product. Interestingly, these modifications may also target the initially added sugar(s), leading to acetylated diglycosides such as scropolioside D, found in different *Scrophularia* species (Ahmed et al. 2003). Oxidation of the iridoid core at carbons 7-8 and formation of a double bond is a key step towards aucubin and catalpol based-iridoids, whereas the persistence of a hydroxyl group on carbon 8 leads to the formation of harpagide and its derivatives (Figure 1, C).

The genus *Scrophularia* (Order Lamiales, Family Scrophulariaceae) includes over 200 figwort species and is particularly studied for its abundance in iridoids and their pharmaceutical activities (de Santos Galíndez et al. 2002; Brownstein et al. 2021). Among these compounds, the most abundant in *Scrophularia nodosa* is harpagoside, with a content of 1% of dry weight (DW) in leaves (Sesterhenn et al. 2007), similar to the concentration found in roots of *H. procumbens* (Order Lamiales, Family Pedaliaceae) (Levielle and Wilson 2002; Mncwangi et al. 2012). The biosynthetic pathway leading to this compound is unclear in both species, but it likely shares its common early *epi*-iridoid part with other species. Early labelling studies in *Scrophularia umbrosa* identified potential key intermediates shared with catalpol and aucubin biosynthesis (Damtoft et al. 1993; Damtoft 1994), but most of the enzymes responsible for carrying the reactions through the pathway to harpagoside have not been identified yet.

From 7-*epi*-deoxyloganic acid, the harpagoside pathway proceeds through several uncharacterized steps, including a decarboxylation at carbon 4, as well as hydroxylations at carbons 5, 6 and 8 required for the formation of harpagide. These reactions are likely catalyzed by cytochromes P450, although the enzymes performing these steps and the precise sequence of the reactions remains speculative (Figure 1, C). The final step of the harpagoside pathway involves the addition of a cinnamoyl moiety on carbon 8 of harpagide. This reaction is expected to be catalyzed by an acyltransferase most likely belonging to the BAHD, SCPL or GDSL families, all of which having been shown to contribute to such acylations in plants. BAHD acyltransferases are named after the first enzymes characterized in this family: benzylalcohol *O*-acetyltransferase (BEAT) (Dudareva et al. 1998), anthocyanin *O*-hydroxycinnamoyltransferase (AHCT) (Fujiwara et al. 1997), anthranilate *N*-hydroxycinnamoyl/benzoyltransferase (HCBT) (Yang et al. 1997) and deacetylvindoline 4-*O*-acetyltransferase (DAT) (St-Pierre et al. 1998). They use acyl-coenzyme A activated donors (Bontpart et al. 2015). Serine carboxypeptidase-like (SCPL) acyltransferase have been less thoroughly studied and use glucose esters as acyl donors (Fraser et al. 2007; Bontpart et al. 2015). Plant GDSL motif-containing esterases/lipases (GDSL) usually perform hydrolase reactions, but some are involved in the synthesis of different classes of mostly parietal metabolites, such as cuticular lipophilic compounds or chlorogenic acid derivatives. GDSL use various acyl donors such as chlorogenic acid or CoA-esters (Yeats et al. 2012; Miguel et al. 2020). No cinnamoyl specific transferase has been discovered to date in any of these families.

In the frame of the study of the harpagoside biosynthetic pathway, we report here the first genome sequencing and assembly of the new iridoid-producing plant model *Scrophularia nodosa*. Using this reference genome together with transcriptomics resources from different plant organs, we identified the enzymes performing the early steps of the iridoid pathway in *S. nodosa.* We then focused our investigations on the last step of the harpagoside pathway and analyzed the features of the large *S. nodosa* BAHD family. We discovered a new group of phylogenetically close sequences in clade 6, clustering together and showing a conserved VYPWG signature motif in the place of the canonical DFGWG motif. In this group, we identified a BAHD enzyme able to catalyze the transfer of a cinnamoyl moiety from cinnamoyl-Coenzyme A to harpagide and to form harpagoside, the main iridoid in *S. nodosa*. More detailed characterization revealed that this enzyme shows a strong specificity towards cinnamoyl-CoA and is unable to use coumaroyl-CoA as an acyl donor, while its acceptor specificity is less stringent. These results open new perspectives in understanding the diversity of reactions carried by BAHD enzymes in specialized metabolism, and in the recombinant production of cinnamoylated compounds.

## Results

### *Scrophularia nodosa* genome and transcriptome

Nanopore genome and cDNA sequencing was performed on young *S. nodosa* leaves tissues from a pseudo-homozygous F7 inbred line, and Illumina RNA-seq was used for differential transcriptome sequencing across various fully developed plant tissues, namely flowers, flower buds, young leaves, older leaves, stems with trichomes (top), stems without trichomes (basal), roots. Quality control PCA analysis of the transcriptomes showed a clustering of samples according to the organ tested, and the main component (PC1 = 64%) appears related to the difference between aerial parts and roots (Supplementary Figure S1). For genome assembly, 99.39% of 708 contigs were assembled into 18 pseudochromosomes (+1 for unplaced contigs) using RagTag (Alonge et al. 2022) and *V. thapsus* genomic data as a reference, as this species was the closest relative sequenced at the time (Badad et al. 2023), resulting in a total genome size of 608.3 Mb. Details about the sequencing results are shown in Figure 2, A. A total of n = 36 chromosomes were observed in haploid cells from flowering buds (Supplementary Figure S2), and analysis of synonymous substitution rates (Ks = 0.4) indicated that an ancient genome duplication or hybridization occurred in *S. nodosa* approximately 30-40 million years ago (Supplementary Figure S2). Sequence annotation revealed 29 653 coding sequences and 64.1 % of repeated sequences, with a total long terminal repeat elements transposons (LTR) content of 51.17 %.

**Figure 2.**
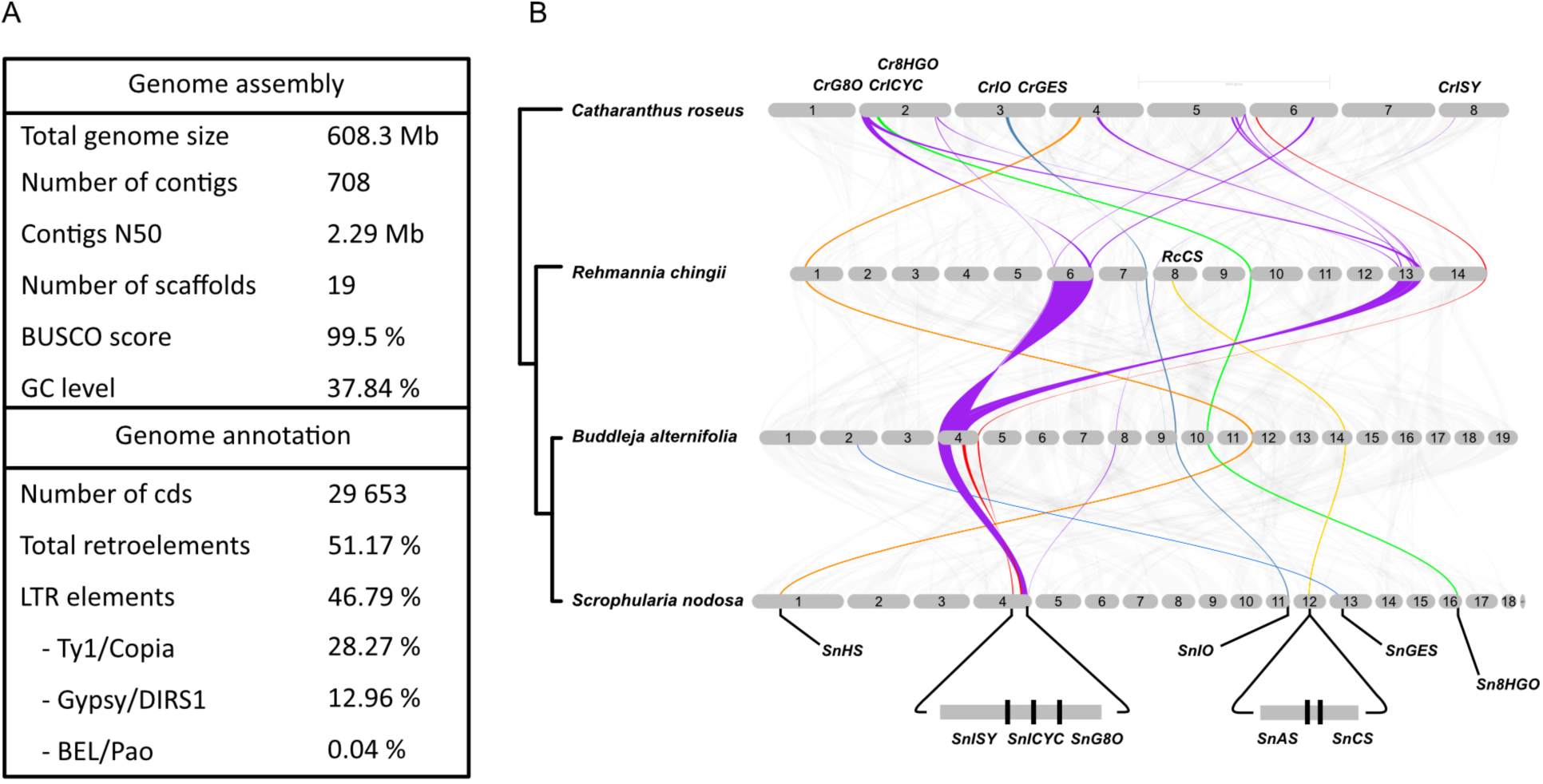
*S. nodosa* genome analysis. **A.** Oxford Nanopore sequencing data *for S. nodosa* gDNA and cDNA. **B.** Block synteny among representative genomes from the Gentianale Apocynaceae (*C. roseus*) and Lamiales Orobanchaceae (*R. chingii*), and Scrophulariaceae (*B. alternifolia*, *S. nodosa*) highlighting the main genes involved in the iridoid pathway, as previously reported for *C. roseus* and *R. chingii*, or discovered in this study for *S. nodosa*. GES: geraniol synthase, G8O: geraniol-8 oxidase, 8-HGO: 8-hydroxygeraniol oxidase, IO: iridoid oxidase, ISY: iridoid synthase, ICYC: iridoid cyclase, CS: catalpol synthase, AS: aucubin synthase, HS: harpagoside synthase.

### *S. nodosa* early iridoid pathway reconstruction

We first identified homologous candidate genes coding for geraniol synthase (SnGES), geraniol 8-hydroxylase (SnG8O), 8-hydroxygeraniol oxidase (Sn8HGO), iridoid oxidase (SnIO), iridoid synthase (SnIS) and 7-*epi*-deoxyloganetic acid glucosyltransferase (Sne7DLGT) in our *S. nodosa* using publicly available data from the related iridoid-producing species *C. roseus*. We found that in our genome assembly, *SnISY*, *SnG8O* and *SnICYC* genes are grouped at the end of scaffold 4, a feature that appears conserved in the other iridoid-producing Scrophulariaceae genome we analyzed (*Buddleja alternifolia*), and to a lesser extent in a representative iridoid-producing Orobanchaceae (*Rehmannia chingii*), but not conserved in the Apocynaceae *C. roseus* (Figure 2, B). Putative homologous genes for the other steps of the early pathway (*SnGES*, *SnHGO, SnIO*) appeared scattered among other assembled *S. nodosa* scaffolds. We then analyzed the gene expression profiles of the candidate genes in our *S. nodosa* RNA-seq dataset in different organs and found them highly correlated (Figure 3, A).

**Figure 3.**
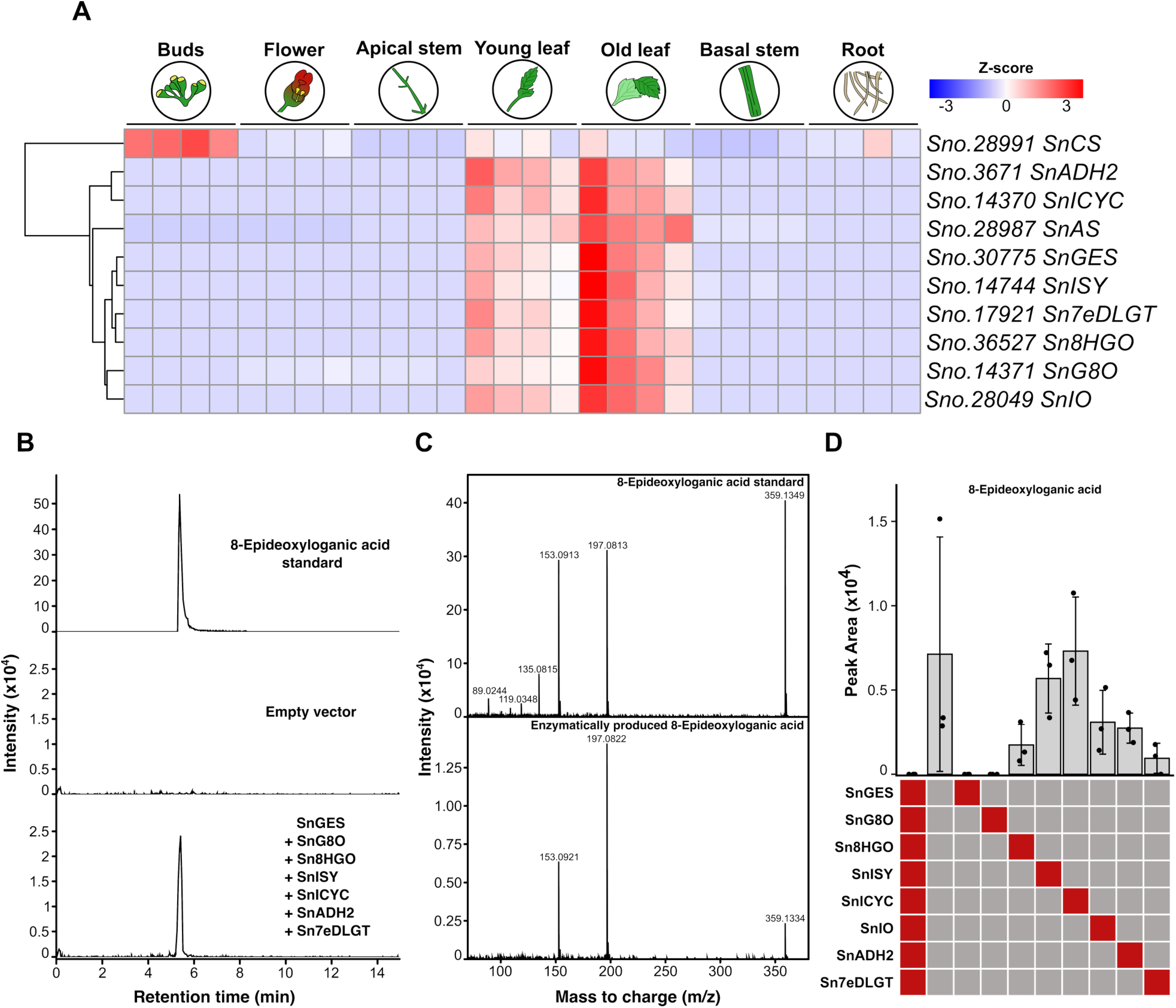
Early iridoid pathway reconstruction. **A.** Hierarchical clustering analysis of expression patterns for early and late iridoid pathway genes in different *S. nodosa* organs, obtained by normalized Z-score of the normalized read counts from DESeq2. 4 replicates are shown for each organ considered. **B.** 8-*epi*-deoxyloganic acid production by transiently expressed enzymes in *N. benthamiana* leaves and comparison with pure compounds (EIC: [M-H]^-^ 359.1348 ± 0.01). **C.** MS2 spectra obtained for 8-*epi-*deoxyloganic acid standard ([M-H]^-^: 359.1353) and enzymatic product ([M-H]^-^: 359.1354). **D.** 8-*epi-*deoxyloganic acid peak area ± standard deviation (n = 3) produced in *N. benthamiana*. Each column shows a combination of genes tested, and red squares indicate omitted gene(s) in the combination.

Next, to proceed with a functional validation, *S. nodosa GES, G8O, 8HGO, IO, IS, 7eDLGT* genes were cloned and “drop-out” combinations, where one of the genes is omitted, were transiently expressed in *N. benthamiana*. Co-expression of all candidates led to the formation of a specific compound when compared to control, with a mass to charge (m/z), retention time and MS/MS fragmentation spectrum corresponding to *epi*-deoxyloganic acid standard (Figure 3, B, C), confirming the functionality of the gene set. Unexpectedly, in our test only *SnGES* and *SnG8O* were essential for the production of *epi*-deoxyloganic acid. Omission of other genes in the combination still allowed the formation of the compound, albeit at lower levels, except for *SnISY* and *SnICYC*, where similar amounts to the full pathway expression were detected (Figure 3D). This result could be explained by the commonly observed spontaneous oxidation and cyclization of the unstable iridodial and iridotrial intermediates. Product formation when *Sne7DLGT* was omitted could be explained by promiscuous glycosylation performed by *N. benthamiana* enzyme(s), as observed previously (Höfer et al. 2013; Ilc et al. 2017).

In the late steps of the iridoid pathways only a few enzymes are known. We identified genes encoding homologs of the recently discovered aucubin synthase (AS) and catalpol synthase (CS) by homology search from the published sequences, in agreement with the fact that *S. nodosa* also accumulates these two molecules. We observed that *S. nodosa AS* (*SnAS*) and CS (*SnCS)* genes are found in close proximity on scaffold 12 of our genome assembly (Figure 2, B). Surprisingly, *SnAS* gene expression correlated with the early pathway down to *Sn7eDLGT*, but *SnCS* expression profile was quite different and more specific to unopened flowering buds (Figure 3, A). Because of the absence of information on the enzyme sequence involved between 7eDLGT and AS/CS in the pathway, we could not hypothesize whether in the harpagoside branch of the pathway the missing genes expression was more likely related to either *Sn7eDLGT/SnAS* or *SnCS* pattern, or plainly different. Given the limited knowledge, commercial availability and low stability of purified compounds for biochemical studies of the central part of the pathway, we started investigating the final step of the pathway, in which harpagide is acylated to form harpagoside, as a starting point for pathway gene discovery.

### Diversification of BAHD sequences in Lamiales

The final step to form harpagoside is the addition of a cinnamoyl moiety on harpagide (Figure 1, C), yet no acyltransferase catalyzing this reaction was described so far. Three enzyme families are reported to contribute to such acylations in plants: SCPL, GDSL lipase-like proteins and BAHDs. A preliminary biochemical assay was performed using semi-purified *S. nodosa* leaves protein extract to evaluate the type of reaction involved in harpagide cinnamoylation. Total soluble proteins were extracted from leaf tissues, precipitated and desalted to remove endogenous harpagoside carry-over. Enzymatic assays revealed a cinnamoyl-CoA dependent harpagoside production from harpagide (Supplementary Figure S3). Consequently, our subsequent efforts focused on the BAHD family members as candidates harpagoside synthase.

As a first step to gather BAHD candidate genes for harpagide cinnamoylation, automated annotation of the *S. nodosa* genome using TMM domain term “transferase” provided a list of 115 BAHD genes. Sequence alignment and phylogenetic analysis showed that *S. nodosa* BAHDs fall into the 7 clades found in terrestrial plants (Figure 4) and defined in recent works (D’Auria 2006; Kruse et al. 2022). *S. nodosa* does not appear to possess significantly more total BAHD genes (68) than the 13 representative iridoid-producing or not producing species whose genome were analyzed (Figure 5, A, B). This number could also be somehow dependent on genome size. We observed an overall high number of BAHD genes in *Nepeta cataria* which could be due to its tetraploid genome (Lichman et al. 2020), yet the diploid *Nepeta racemosa* shows a similar number. As for the higher number observed in *Scrophularia ningpoensis*, although behaving as a diploid species, this close relative to *S. nodosa* has been shown to possess n = 72 chromosomes (Xu et al. 2025), *i.e.* twice the number we observed in our *S. nodosa* samples. Clade 6 BAHDs are reported to include enzymes active on terpenoids and alkaloids (Moghe et al. 2023) and were further examined. Within this clade, *S. nodosa* again did not display an unusually high number of genes (22) compared with other species (Figure 5, C). Based on sequence similarity and previously defined subclades (Kruse et al. 2022), clade 6 was annotated into 5 sub-clades (a, c, d, h and i). Of the 22 *S. nodosa* sequences, 4 grouped in sub-clade 6c, 5 in sub-clade 6h, and 13 in sub-clade 6i (Figure 5, C).

**Figure 4.**
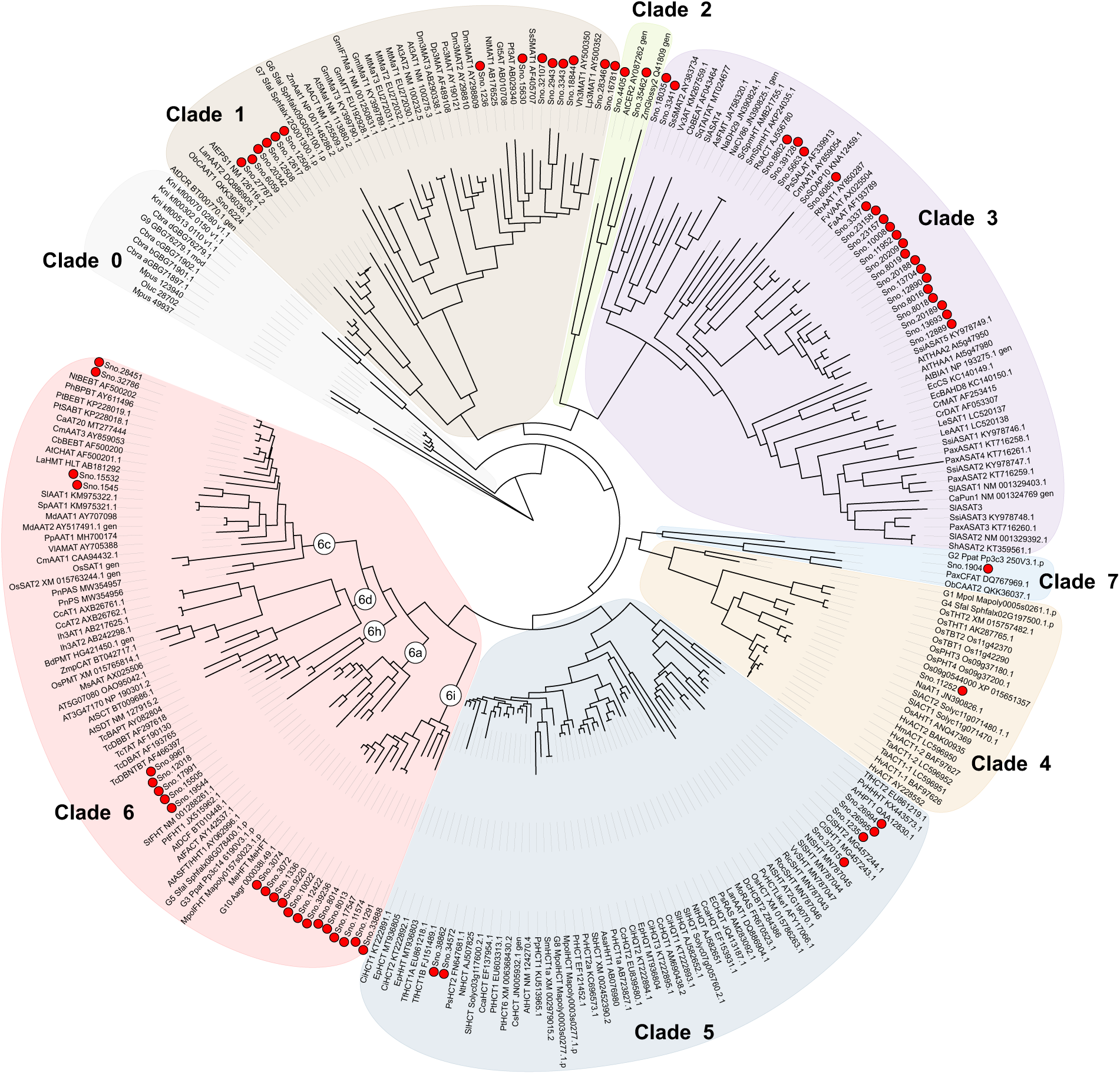
Phylogenetic tree of *S. nodosa* BAHD and 46 reference plant BAHD amino acids sequences (Kruse et al. 2022). Clades and clade 6 subclades are indicated. Red dots indicate *S. nodosa* sequences.

From this analysis, we observed that the genomes of Lamiales are particularly enriched in sub-clade 6i BAHDs sequences (Figure 5, D). At the base of this subclade, a number of proteins (but none from the Scrophulariaceae family) retain the canonical DFG(F/W)G motif characteristic of BAHD enzymes. However, a significant sequence divergence occurred within subclade 6i, giving rise to the formation of 3 separate branches characterized by a more hydrophobic VYPWG motif (branches 1, 2, 3 in Figure 5, D, E). Among these, branch 2 contains exclusively Lamiales sequences (including 13 from *S. nodosa*), and branch 3 contains only Scrophulariaceae sequences including most (9) of clade 6 *S. nodosa* BAHDs (Figure 5, E). The specific occurrence of branches 2 and 3 in Lamiales and Scrophulariaceae could thus be linked to their specialized terpenoid metabolism. This prompted us to consider as best candidates the 13 *S. nodosa* sequences from the clade 6i, in particular those of branches 2 and 3.

### Gene expression analysis identifies an harpagoside synthase candidate gene

To further support and narrow-down our BAHD candidate selection, we then compared the expression profiles of the 21 *S. nodosa* clade 6 BAHD-encoding genes (with expression read counts above 500) to harpagoside concentration in 6 different plant tissues using Pearson correlation (Figure 7, A). *S. nodosa* young leaves and unopened flowering buds were the main harpagoside accumulating organs, with approximatively 5 mg / g fresh weight (FW), while older leaves appeared to contain much less harpagoside (approx. 0.8 mg / g FW) (Figure 7, B). *Sno.1336* was the highest ranked gene in our correlation analysis with a Pearson correlation coefficient (PCC) of 0.84, showing a high global level of expression, mostly in unopened flowering buds and younger leaves compared to the other tissues tested (Figure 7, A). *Sno.1336* was thus selected as a prime candidate for recombinant expression and enzymatic studies.

### Biochemical characterization of *S. nodosa* harpagoside synthase

*Sno.1336* cDNA was cloned into the pEPStrepGW vector and expressed in *Komagataella phaffii* (*Pichia pastoris*) GS115. The Sno.1336 enzyme was purified from the recombinant yeast by Strep-tag affinity chromatography. Protein activity was then tested using different acyl-CoA donors and iridoid acceptor molecules. Sno.1336 efficiently catalyzed the formation of harpagoside from harpagide and cinnamoyl-CoA (Figure 7, C, E). The reaction product was confirmed using high resolution LC-MS/MS and harpagoside standard (Figure 7, D). Accordingly, Sno.1336 was named *S. nodosa* harpagoside synthase (SnHS).

Using different acyl acceptors, substrate preference of SnHS was investigated. It showed the highest activity with harpagide and also ajugol, the supposed direct precursor of harpagide in the pathway and differing only from harpagide by the absence of hydroxylation at carbon 5 (Figure 1, C; Figure 8, A). Loganic acid was also converted to a cinnamoylated product, although this compound is not present in the harpagoside pathway. Regarding acyl donors used by SnHS, cinnamoyl-CoA was by far the most efficient physiologically relevant substrate (*Km* = 192 µM), but synthetic *ortho*-coumaroyl-CoA was also used with comparable efficiency in an end-point assay (Figure 8, B). In contrast, only very low conversion was measured using the physiologically relevant *para*-coumaroyl-CoA.

## Discussion

Iridoids large spectrum of biological activities (hepatoprotective, antimicrobial, anti-inflammatory, anticancer, among others) have been known for centuries through the traditional use of many plant species, in particular in the Lamiaceae and Scrophulariaceae families (Tundis et al. 2008; A. Viljoen et al. 2012; Mncwangi et al. 2012). Iridoid-producing species thus constitute a vast reservoir of valuable molecules and scaffolds for drug design, and the global interest for these molecules is evidenced by the exponentially increasing number of studies on the topic. The number of publications on “plants” increased by a factor of 10 from 1990 to March 2026, while in the same time the number of articles citing the term “iridoid” in NCBI PubMed database has grown from 220 to over 7000. These studies mainly concentrated on isolation and identification of new iridoids, notably through purification and NMR, and plant extract bioactivity screening (see (Dinda et al. 2007b, 2007a, 2009, 2011) for extensive review).

Owing to the recent development of fast and inexpensive gDNA and RNA sequencing and analysis techniques and to the optimization of recombinant expression systems such as transient expression in *N. benthamiana*, the diversity of iridoid-producing species can now be explored without much constraints and their biosynthetic pathways deciphered. Focusing our interest on harpagoside, an anti-inflammatory iridoid found in a few angiosperm species (Georgiev et al. 2013; Brownstein et al. 2017), we aimed at the discovery of the specific enzyme set needed to build its structure. *H. procumbens*, the first species shown to accumulate harpagoside and one largely used in phytotherapy, is an endangered desert plant from southern Africa, accumulating harpagoside mostly in its roots (Levieille and Wilson 2002). The destructive methods of root harvesting and very slow growth of the plant in its natural habitat are inherently not sustainable, especially with an increasing global demand in harpagoside for health applications in human and animals. Protection and improved harvesting practices measures, as well as *in vitro* micro-propagation have been implemented to preserve *H. procumbens* wild populations (Stewart and Cole 2005). The species is also amenable to callus and plant regeneration, but the rates of harpagoside production are still low to not detectable (Sesterhenn et al. 2007; Grąbkowska et al. 2014, 2016). We propose an alternative model species suitable for both biological studies and harpagoside production. *S. nodosa*, a perennial figwort growing commonly in the wild in Western Europe, shows both promising harpagoside accumulation (approx. 5 mg / g FW in leaves), ease of greenhouse cultivation in standard conditions, and a life cycle amenable to 2-3 months in controlled conditions. Contrary to *H. procumbens*, *S. nodosa* contains significant harpagoside concentrations throughout the plant and especially in leaves, allowing its potential non-destructive harvesting. The elucidation of the *S. nodosa* harpagoside biosynthetic pathway, starting with the acquisition of sufficient knowledge about its genome and genes involved, is thus a first crucial step towards optimization and engineering of harpagoside production in plants or in recombinant organisms.

Nanopore sequencing revealed that *S. nodosa* possesses a medium size, diploid genome, which we assembled into 19 scaffolds. Gene sequences and gene expression profiles obtained from Illumina sequencing of different plant organs allowed us to identify functional homologs for the early iridoid pathway leading to the biosynthesis of e7DLGA, and we achieved successful recombinant reconstruction of this early part of the pathway in *N. benthamiana.* Sequence analysis showed that the enzymes involved in the early iridoid pathway are highly conserved between *S. nodosa* compared with the model species *C. roseus*, although the two species belong to two different orders, Lamiales and Gentianales, respectively. Despite this conserved function, major differences in the physical gene organization of the early pathway are detected between the two species. In *C. roseus*, *G8O*, *ICYC* and *8HGO* are clustered on chromosome 2, while *ISY* is located on chromosome 6. In *S. nodosa*, *G8O*, *ISY* and *ICYC* are clustered at the extremity of scaffold 4, whereas *8HGO* is found on scaffold 16.

We also found *S. nodosa* aucubin synthase (*SnAS*) and catalpol synthase (*SnCS*) homologs present as neighbor genes on scaffold 12, and nonetheless very differently regulated in terms of expression, with *SnAS* following the same pattern as the early pathway genes, while *SnCS* being mainly expressed in unopened flower buds. This is different from the coregulation regularly observed for clustered genes involved in specialized metabolic pathways (Shan et al. 2025), and might underline a particular need for differential tissue accumulation of aucubin and catalpol with possibly a transport of aucubin from the young leaves to the flower buds. Aucubin is indeed not a mere transient precursor of catalpol, but is a significantly accumulated metabolite both in *S. nodosa* and other iridoid-producing species (Rønsted et al. 2000; Xie et al. 2020). Reports of detection and isolation of numerous iridoids derived from harpagide, bartsioside, ajugol (Dinda et al. 2007a, 2007b, 2009, 2011) may indicate that late iridoid pathways are not straight lines towards final bioactive molecules (e.g. catalpol, harpagoside), but are the just the longest branches along which intermediates can accumulate and/or be further decorated according to the species, organ, or growth/ecological conditions considered.

In the first step of our study, we functionally validated a conserved chassis for the early pathway in *S. nodosa*. It will be of significant help in the further investigate the iridoid pathways. The central section of the pathway being more complex to characterize due to instability and unavailability of the metabolic intermediates, our next priority was to identify the enzyme catalyzing final step of the pathway, the cinnamoylation of harpagide into harpagoside. The combination of phylogenetic analysis, correlation between gene (co)-expression and enzyme activity, led to identification of a candidate protein that could be demonstrated *in vitro* to catalyze harpagoside formation from cinnamoyl-CoA and harpagide, the most abundant substrate in leaves (Supplemental Figure S5), and was named harpagoside synthase (HS). This first cinnamoyl transferase characterized in plants shows specificity for a few iridoid structures and cinnamoyl-CoA and *O*-coumaroyl-CoA, both unsubstituted on the 4 ring position, and belongs to the family of the BAHD acyltransferases.

A phylogenetic analysis of the BAHD proteins revealed the evolution of specific branches of clade 6 within Lamiales and more specifically Scrophulariaceae (branches 2 and 3 in clade 6i) including this new HS, characterized by an atypical VYPWG motif instead of the typically conserved DFGWG motif found in most BAHDs (Figure 6, E). The latter motif has been shown to locate on the protein surface and far from the catalytic HxxxD dyad (Supplemental Figure S6, B), but site-directed mutagenesis studies have shown from a slight reduction in activity to activity loss was observed when the fist Asp residue of the DFGWG motif was replaced in vinorine synthase from *Rauvolfia serpentina* and anthocyanin 5-*O*-glucoside-6-*O*-malonyltransferase from *Salvia splendens* (Suzuki et al. 2003; Bayer et al. 2004). Based on enzyme structure (Ma et al. 2005), this motif is expected to have an important structural role in the organization of the loop between two alpha-helix domains. A role in the CoA ester access or channeling cannot be excluded. The increasing number of available BAHD sequences revealed other variations in this DFGWG motif. A particularly studied example is clade 2 enzymes involved in long-chain fatty acids machinery and with no BAHD activity known to date. The DFGWG motif is not conserved in this clade and replaced by EIN/KGG/Q in *Arabidopsis thaliana* AtCER2 and maize ZmGlossy2 (Supplemental Figure S4). More investigation will be needed to test a possible link between the DFGWG shift to VYPWG sequence, and selectivity in acyl donor or acceptor in BAHD clade 6 enzymes, as the observed sequence diversification in the branch 3 of clade 6i is paralleled with the occurrence of a significant number of cinnamoylated compounds in Scrophulariaceae (Pasdaran and Hamedi 2017). Interestingly, no sequence belonging to this branch was found in the genome of *Sesamum indicum*, a representative of the Pedaliaceae family which also includes *H. procumbens*, the current main source of harpagoside for medical applications. This may suggest that the formation of harpagoside in *H. procumbens* may result from a convergent evolution.

**Figure 6.**
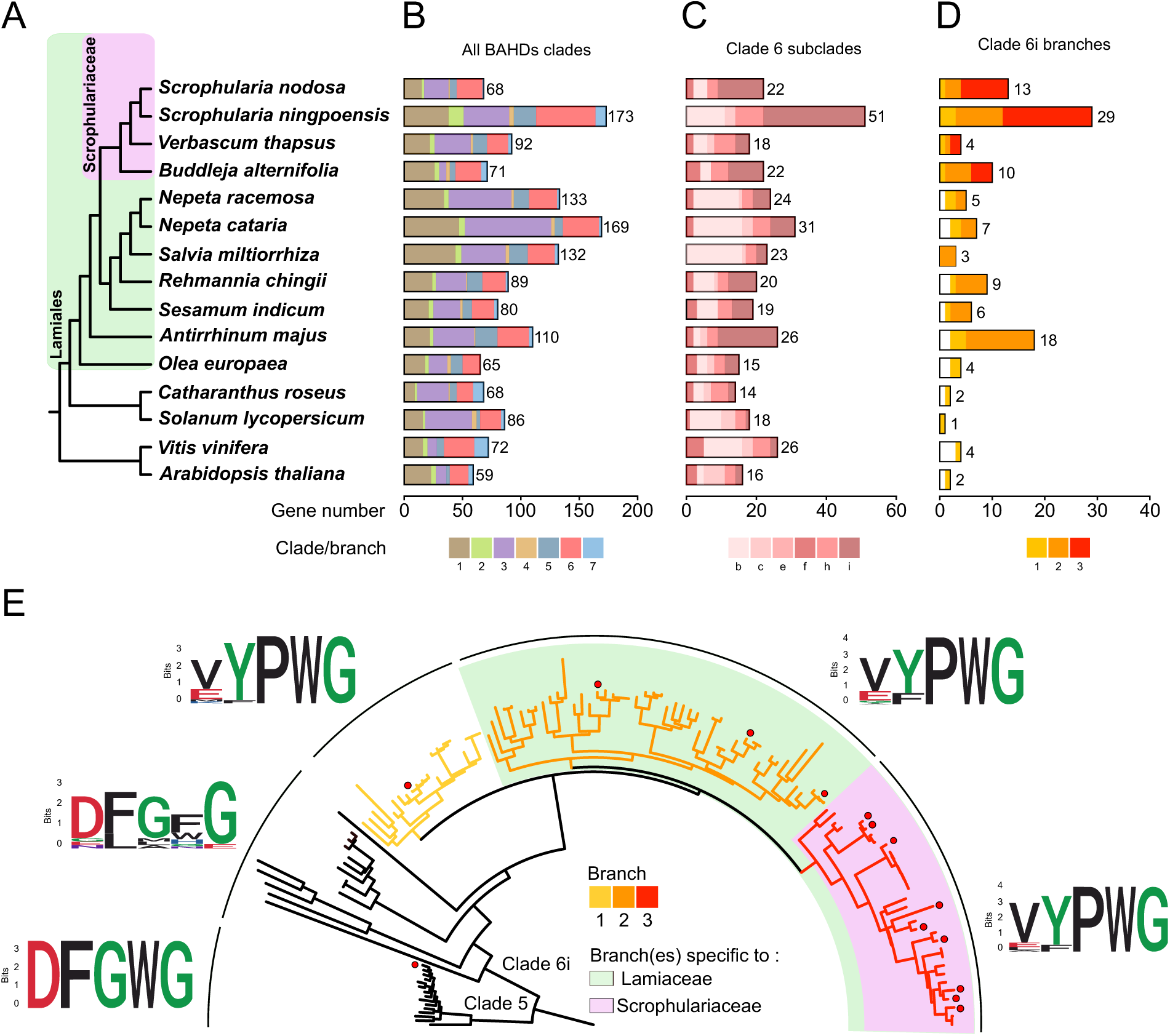
Analysis of the *S. nodosa* BAHD family protein sequences. **A**. Cladogram of the different species examined. **B.** Total number and clade repartition of total BAHD genes identified by HMM motif “transferase”. **C**. Total number and repartition into different subclades of clade 6 BAHD genes. **D.** Total number and specific branches (1, 2, 3) of subclade 6i BAHD genes. **E.** Phylogenetic tree of subclade 6i BAHD protein sequences and analysis of the conserved “DFGWG” motif per branch. Clade 5 representative sequences were used to root the tree. Red dots indicate *S. nodosa* sequences. Consensus (Weblogo) sequences of the canonical “DFGWG” motif found in the different branches are displayed; acidic amino acids are shown in red, basic in blue, hydrophobic in black, neutral in purple and polar in green. Detailed phylogenetic tree is shown in Supplemental Figure S6.

**Figure 7.**
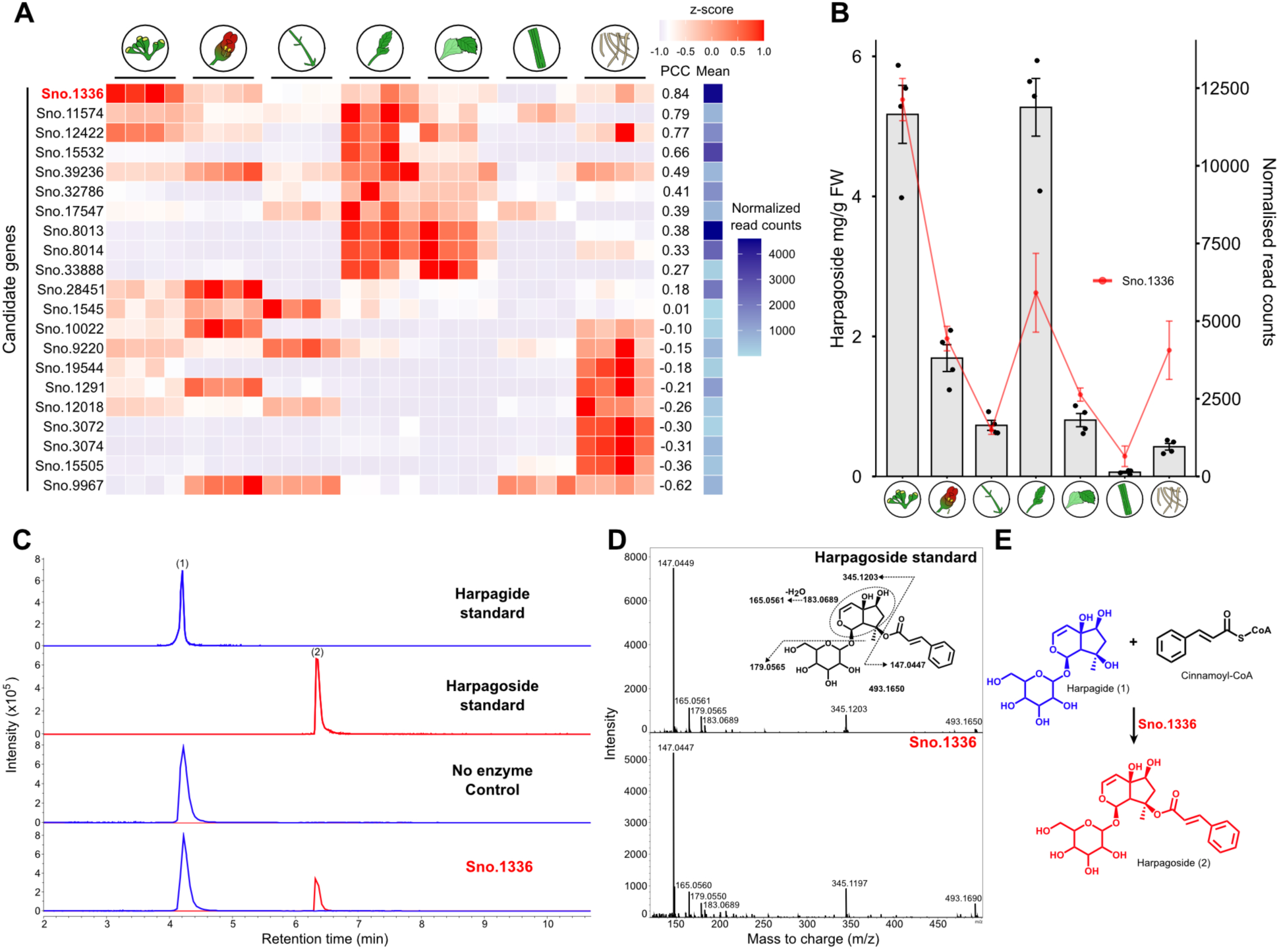
Validation of the harpagoside synthase activity of Sno. 1336. **A**. Correlation between harpagoside content and clade 6 BAHD genes expression in different *S. nodosa* organs (from left to right: unopened buds, flowers, trichome-bearing stems, young leaves, mature leaves, basal stems without trichomes, roots, 4 biological replicates per organ are shown). Genes are ordered according to the Pearson correlation coefficient (PCC) relative to harpagoside content. Global expression level of each gene is shown as the mean of normalized read counts. **B**. Harpagoside content (mg/g fresh weight) and *Sno.1336* expression in different plant tissues. **C.** Sno.1336 harpagoside synthase activity detected by conversion of harpagide (EIC: [M-H]^−^: 363.1297 ± 0.01; [M+HCOOH-H]^−^ = 409.1351 ± 0.01) and cinnamoyl-CoA to harpagoside (EIC: [M-H]^−^: 493.1715 ± 0.01; [M+HCOOH-H]^−^ = 539.1770 ± 0.01) and comparison with pure compounds. **D.** MS2 spectra obtained for the harpagoside standard ([M-H]^−^: 493.1715) and the Sno.1336 product ([M-H]^−^: 493.1716). **E.** Details of the common fragments mapped onto the harpagoside structure.

**Figure 7.**
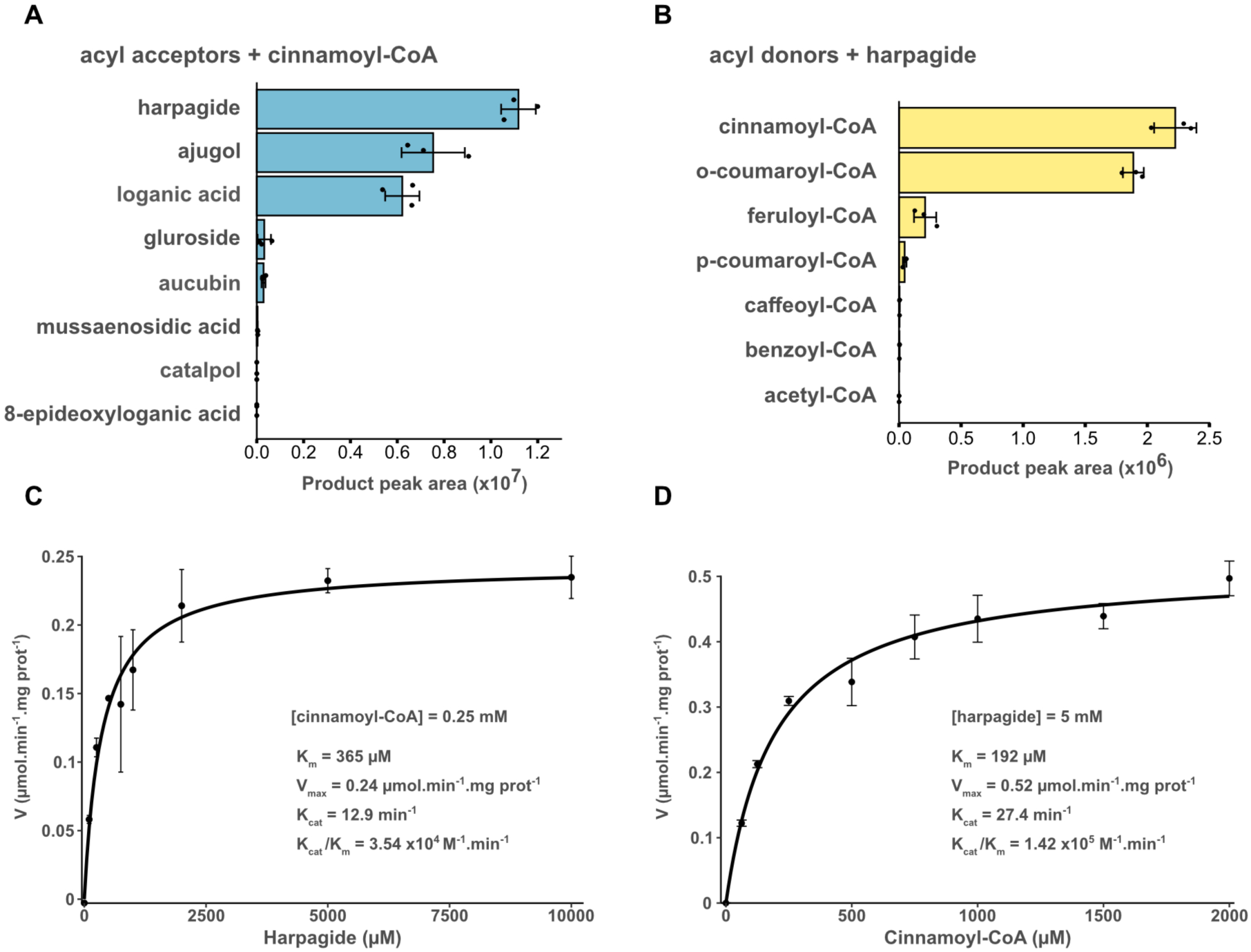
Biochemical characterization of *S. nodosa* harpagoside synthase Sno 1336. **A.** *In vitro* conversion of different acyl acceptors (2.5 mM) using cinnamoyl-CoA (0.5 mM) as acyl donor. **B**. *In vitro* conversion of different acyl donors (0.5 mM) using harpagide (2 mM) as acceptor. **C, D.** *K_m_* and *V_max_* determination for the two main substrates of SnHS, harpagide and cinnamoyl-CoA.

We established here *Scrophularia nodosa* as a new model to explore iridoid metabolism and harpagoside production. Our approach demonstrates that unknown steps of biosynthetic pathways can be deciphered using a combination of high-quality gDNA assembly and differential RNA-seq data, together with a rational selection of candidates for investigation via crude extract activity pre-screening and detailed protein sequence analysis. The use of different complementary screening methods has proven essential to analyze complex and highly variable enzyme families such as BAHDs. This workflow will form the basis of future studies on the complex iridoid diversity in our model, *S. nodosa*, and provides a unique tool for the production of cinnamoylated compounds.

## Material and Methods

### Chemicals

Harpagide, harpagoside, aucubin, catalpol were purchased from Extrasynthèse (Genay, France). Cinnamoyl-CoA, ortho- and para-coumaroyl-CoA, benzoyl-CoA, acetyl-CoA, feruloyl-CoA, caffeoyl-CoA were purchased from AnalytiCon Discovery (Potsdam, Germany).

### Plant material

*S. nodosa* seeds were purchased from Graines de vie (Dontreix, France), and were germinated and cultivated at 25°C, 16h hour photoperiod. One plant was used for self-fertilization at each of 7 cultivation cycles. 7^th^ generation plants were harvested when flowering was advanced and parts/organs were sampled: flowers, flower buds, apical trichome-bearing stems, basal trichome-free stems, apical young leaves, basal older leaves, roots. 6 replicates were collected, each formed by pooling tissues from 3 plants, ground in liquid nitrogen and stored at -80°C.

### Oxford Nanopore DNA sequencing

Genomic DNA was isolated from young leaves of *Scrophularia nodosa* using a Nucleon Phytopure Genomic DNA Extraction Kit (Cytiva) based protocol followed by size selection with the SRE-XL size selection kit (Circulomics). DNA quality was checked on a 1% agarose gel. Absorption ratios A_260nm_/A_280nm_ and A_260nm_/A_230nm_ were measured on a Nanodrop 2000 spectrophotometer (ThermoFischer Scientific) and accurate quantification was done with the Qubit DNA HS assay kit (Fischer Scientific). DNA library was prepared using the SQK-LSK114 kit (Oxford Nanopore Technologies). The final library was loaded on a PromethION R10.4.1 flowcell (Oxford Nanopore Technologies) and sequenced with the PromethION P2solo device (Oxford Nanopore Technologies).

### Illumina RNA sequencing

Total RNA was extracted from fresh tissue powder using Concert Plant RNA Reagent (Invitrogen) according to the manufacturer’s instructions. RNA samples were treated with DNAse I (Thermo Fisher Scientific) followed by phenol/chloroform extraction. RNAs concentrations were determined using a QuBit Fluorometer (Invitrogen). RNA integrity was checked using a 2100 Bioanalyzer (Agilent). Libraries were prepared using the NEBNext® Poly (A) mRNA Magnetic Isolation Module (NEB) for mRNA isolation combined with the NEBNext Ultra II Directional RNA Library prep kit for Illumina (NEB). The libraries were sequenced using an Illumina Nextseq 2000 system (single-end mode 1 × 50 bp) for 9 conditions (different tissues) in quadruplicates (x4).

### Oxford Nanopore cDNA library preparation & sequencing

cDNA library was prepared using the SQK-PCR109 kit (Oxford Nanopore Technologies) with 50 ng of total RNA as starting material. The final library was checked on the Bioanalyzer 2100 & High Sensitivity DNA Assay (Agilent) and loaded on a PromethION R9.4.1 flowcell (Oxford Nanopore Technologies). Sequencing was done with the PromethION P2solo device (Oxford Nanopore Technologies).

### Genome and transcriptome analysis

Nanopore transcriptome cDNA sequencing generated 30.8 Gb of raw data. Basecalling was performed using dorado v0.7.2 with model *dna_r9.4.1_e8_sup@v3.6*, and out reads smaller than 100bp were filtered out to finally obtain 26.5 Gb of data.

Nanopore DNA-seq sequencing was performed using two PromethION2 R10 flowcells with 49.9 Gb of data. Basecalling was performed using dorado v0.7.2 with model *dna_r10.4.1_e8.2_400bps_sup@v4.1.0*, and reads shorter than 1kb were filtered out for a total of 41.8 Gb of data. All nanopore raw files (.pod5) were basecalled in “super-accuracy” mode with dorado.

### Genome *de novo* assembly

*De novo* assembly was performed using NextDenovo (v2.5.2) with read correction option (task = all). The assembly resulted in a total of 708 contigs with an N50 of 2.29 Mb and a total genome size of 608 Mb. Assembly completeness was assessed using BUSCO (v5.7.1) and the *viridiplantae_odb10* database, with a score of 99.5% [S:83.3%,D:16.2%]. The obtained contigs were assembled into chromosome-level scaffolds by making use of RagTag (v2.1.0) in scaffold mode. *Verbascum thapsus* (daVerThap1.1) was used as reference genome to set the number of chromosomes as the closest species with a completed genome at the time of analysis. Unplaced contigs were grouped into a single scaffold called “ChrUn”.

### Transcriptome *de novo* assembly

Nanopore cDNA reads were aligned using minimap2 v2.26 (with options -ax splice -G 4k) and NT paired-end Illumina RNA-seq data were aligned with hisat2 (with option --max-intronlen 4000) to our new assembled genome. The cDNA and RNA-seq alignments were fed to StringTie2 v2.2.0 (--mix option) to identify transcripts and their isoforms. Stringtie2’s list of transcripts was then passed to Transdecoder (v5.7.1) for ORFs prediction. We obtain a total of 71 527 isoforms for 29 653 predicted genes. Functional annotation was performed by protein homology search using BLASTp against the SwissProt database, protein family classification using Pfam and HMMER (hmmscan) and signal peptide prediction using SignalP.

### RNA-seq and Differential Gene Expression Analysis

Illumina RNA-seq libraries were mapped using hisat2 v2.2.1 and read counts were quantified for the 29 653 predicted genes using featureCounts v2.0.6. Differential gene expression analysis was conducted in R using the DESeq2 package.

### Cloning and recombinant protein expression

*Sno.1336* was amplified by PCR from *S. nodosa* leaf cDNA and the PCR product was then purified using magnetic beads to remove primers. Gateway technology (Invitrogen) was used to transfer the purified PCR product into the pDONR207 entry vector and finally into the pEPStrepGW expression vector (modified from pPICHOLI). The expression vector was then introduced into *K. phaffii* (*P. pastoris*) GS115 as described in pPiCHOLI handbook. A preculture was grown overnight in YPD medium supplemented with 100 μg.mL^-1^ of Zeocin (InvivoGen) at 30 °C with shaking. BMMY (Buffered Methanol-complex Medium) medium supplemented with 100 μg.mL^-1^ of Zeocin was then inoculated with the overnight culture (10% final concentration of cell suspension) and grown at 30°C with shaking until OD600 reached 1.0. Protein expression was then induced with the addition of 1% methanol (v/v) twice a day (morning and evening) and grown at 30°C with shaking for 2-3 days. The cells were then harvested by centrifugation at 2000 g at 4 °C for 10 min and stored at -20 °C until purification.

For protein purification, cells were resuspended in 100 mM Tris-HCl buffer, pH 8, supplemented with 150 mM NaCl, 1 mM EDTA, and 5 mM dithiothreitol then lysed during 7 cycles at 1700 bars using LM20-30 microfluidizer. The lysate was then centrifuged at 15,000 g at 4 °C for 20 min. Soluble proteins were then loaded onto a Strep-Trap HP column (Cytiva) using an Äkta Pure 10 protein purification system (Cytiva). The recombinant protein was eluted using 100 mM Tris-HCl, pH 8, supplemented with 150 mM NaCl, 1 mM EDTA, and 5 mM desthiobiotin. Fractions containing the protein of interest were then loaded onto a HiTrap Desalting column and eluted using 100 mM Tris-HCl pH 8. Glycerol (10%) was added into fractions containing the protein before storage at -20°C. Concentration of total protein was measured using Qubit Fluorometer (Thermo-Fisher) according to the manufacturer’s protocol. The putative size of the protein was determined via western-blot and then stained with Coomassie blue.

### Biochemical characterization

Stock solution of acceptor substrates and donor acyl-coA donor were prepared in H_2_O. *In vitro* assays were performed in 100 µL containing 50 mM Tris-HCL pH 7.4 using 500 ng of recombinant protein. Reactions were initiated by the addition of enzyme, incubated at 30°C in a water bath for 1 hour and stopped by adding of 100 µL acetonitrile. To determine kinetic parameters, substrate concentrations were optimized for each tested substrate. For harpagide, 0.250 mM of cinnamoyl-CoA and 0.1-10 mM of harpagide were used. For cinnamoyl-CoA, 5 mM of harpagide and 0.0625-2 mM of cinnamoyl-CoA were used. Harpagoside formation was then quantified by HPLC-UV at 280 nm. Kinetics curves and parameters were then calculated using RStudio software.

To investigate the substrate preference, eight iridoid glycosides (ajugol, harpagide, loganic acid, gluroside, aucubin, mussaenosidic acid, catalpol, 8-epideoxyloganic acid) were used as acceptors and seven acyl-CoA (cinnamoyl-CoA, ortho- and para-coumaroyl-CoA, benzoyl-CoA, acetyl-CoA, feruloyl-CoA, caffeoyl-CoA) were used as donors. For acceptors, 0.5 mM of cinnamoyl-CoA and 2.5 mM of iridoid acceptors were used. For acyl donors, 2 mM of harpagide and 0.500 mM acyl-CoA donor were used. Product intensity was then monitored by UHPLC-qToF, putative product EIC was traced based on their chemical formula and peak area intensity measured.

### Early iridoid pathway reconstruction in *N. benthamiana*

Candidate genes sequences were amplified by PCR from *S. nodosa* cDNA and were purified using magnetic beads to remove primers. Gateway technology (Invitrogen) was used to transfer the purified PCR product into the pDONR207 entry vector and finally into the pH2GW7 expression vector. *A. tumefaciens* GV3101 cells were transformed through electroporation (25 µF, 2,5 kV, 400 Ω), recovered in LB without antibiotics at 28°C during 3h and incubated on LB plates containing antibiotics (rifampicin, gentamycin and spectinomycin) at 28°C for 48 h. Agrobacteria were grown overnight in liquid LB medium containing antibiotics (rifampicin, gentamycin and spectinomycin) at 28°C 260 rpm. Cells were harvested by centrifugation at 2000g, 4°C during 15 min and resuspended in infiltration buffer (10 mM MES, 10 mM MgCl2, 100 μM acetosyringone, pH 5.6). OD_600_ was measured and strains were mixed and diluted in infiltration buffer to OD_600_ = 0.1 per strain. A strain harboring the p19 gene was co-infiltrated in all cases. After incubation at 28°C for 2-4h in buffer, mixtures were infiltrated into 4 weeks old *N. benthamiana* leaves and plants grown for 3 days (in greenhouse at 25°C with 16h photoperiod). Leaf material was collected by flash freezing in tubes containing glass beads. 0.1 mg of leaf material was grounded using Precellys Evolution (Bertin) using the soft tissue setup (10000 rpm, 2*30s with 25s hold). Samples were extracted with methanol (1:10, m/v) at 1400 rpm, 4°C overnight using Thermomixer Comfort (Eppendorf). Extracts were then centrifugated at 15000 g for 20 min at 4°C and resulting supernatants were transferred into glass vials. Extracts were then injected in UPLC-QTOF.

### UPLC-QTOF analysis

Sample analysis was performed on an UltiMate 3000 system (Thermo Fisher Scientific) coupled to an Impact II (Bruker) QTOF spectrometer. Chromatographic separation was performed on an Acquity UPLC HSST3 C18 column (2.1 × 100 mm, 1.8 μm; Waters) and using a gradient of solvents A (water, 0.1% formic acid) and B (acetonitrile and 0.1% formic acid). Chromatography was carried out at 35°C with a flux of 0.3 ml/min, starting with 5% B for 2 min and reaching 100% B at 10 min, holding 100% for 3 min and coming back to the initial condition of 5% B in 2 min (15 min). Ionization parameters of ESI source were as follow: acquisition was done in negative ionization mode on a 75 to 1000 m/z mass range with a spectrum rate of 8 Hz and using the AutoMS/MS fragmentation mode. The end plate offset was set at 500 V, capillary voltage at 4500 V, nebulizer at 2 bar, dry gas at 8 liters/min, and dry temperature at 200°C. The transfer time was set at 40.8 to 143 μs and MS/MS collision energy at 50 to 100% with a timing of 50 to 50% for both parameters. The MS/MS cycle time was set to 2 s, absolute threshold was set to 816 counts per second (cts), active exclusion was used with an exclusion threshold at three spectra, released after 1 min, and an ion was reconsidered as precursor for the fragmentation if the ratio current intensity/previous intensity was higher than 5. MS/MS collision energy was set according to the mass, ranging from 6 eV to 35 eV.

### Metabolite extraction from *S. nodosa* tissues

The same plant samples were used both for RNA sequencing and harpagoside quantification. Briefly, *S. nodosa* flowers, flower buds, apical trichome-bearing stems, basal trichome-free stems, apical young leaves, basal older leaves and roots were sampled on fully grown plants. Samples were flash-frozen in liquid nitrogen, ground and stored at -80°C until extraction. 50-100 mg of fresh material was extracted with methanol (1:10, v/v) at 1400 rpm at 20°C for 1 hour using Thermomixer Comfort (Eppendorf). Extracts were centrifugated at 14000 g for 20 min at 4°C and resulting supernatants were transferred into glass vials. Some samples were diluted to avoid detector saturation and peak areas were corrected by corresponding dilution factors.

### HPLC-UV harpagoside quantification

Harpagoside quantification was done on a high-performance liquid chromatography system (Alliance 2695; Waters) coupled to a photodiode array detector (PDA 2996; Waters). 10 µL of sample was injected into Kinetex Core-Shell C18 column (4.6 x 150 mm, 5 µm; Phenomenex). Needle and injection loops were successively washed with weak (95% water/5% acetonitrile) and strong (20% water/80% acetonitrile) solvent mix. The mobile phase consisted of a mix of HPLC grade water (A) and acetonitrile (B), both containing 0.1% formic acid. The elution program was as follows: from 5% B at 0.0 min to 95% B at 13.0 min, hold at 95% B for 2 min and then going back to initial conditions with 5% B in 2 min (17 min). The flow was set to 1 mL.min^-1^, column temperature to 35°C and the absorbance was recorded between 220 and 400 nm. Harpagoside quantity was monitored at 280 nm and data were processed with the Empower 3 Software (Waters).

### Phylogenetic analysis

Available data from *A. thaliana*, *C. roseus*, *S. lycopersicum*, *V. vinifera*, *O. europeae*, *A. majus*, *S. indicum*, *N. cataria*, *N. racemosa*, *R. chingii*, *S. miltiorrhiza*, *B. alternifolia*, *V. thapsus*, *S. ningpoensis* were subjected to protein family classification using pfam and HMMER (hmmscan) as for *S. nodosa.* Protein sequences classified as transferase (PF02458) using hmmscan were used for further phylogenetic analysis. Sequences below 400 amino acids were removed from the dataset. Previous classification (Kruse et al. 2022) was used to sort BAHD sequences from each species into clades. Sequences were then aligned using muscle 5.3 algorithm (Edgar 2022). Maximum likehood phylogenies were constructed using iqtree v2.3.6 using the following parameters -alrt 1000 -B 1000 - nt auto. Phylogenetic trees were visualized and edited using iTOL (https://itol.embl.de) and then annotated using Inkscape. BAHD counts were formatted with RStudio.

## Acknowledgments

We thank Dr Anne Molitor, Antoine Hanauer and Dr Raphael Carapito from the GENOMAX platform of INSERM UMR S1109 for Next-Generation Sequencing experiments. The authors acknowledge the Gene Expression Analysis, p3P, Plant production and MASS facilities at IBMP for their help acquiring the data presented. The authors are grateful to Dr Alexandre Berr for *S. nodosa* chromosome images, and to Prof. Dr. John C D’Auria for providing the pEPStrepGW vector.

## Author Contributions

Experimental design: DR, EGa, DW, NN. Manuscript writing and edition: DR, SW, EGa, DW, NN. Experimental work: DR, SW, AP, YL, DP, CP, EGr, LM, AA, SK.

## Data availability

*S. nodosa* genome, RNA-Seq data, gene sequences and associated data will be available in public databases upon publication (NCBI, Zenodo).

## Supplementary Data files

Supplementary Figures 1 to 6

## Supplemental figures

**Figure S1.**
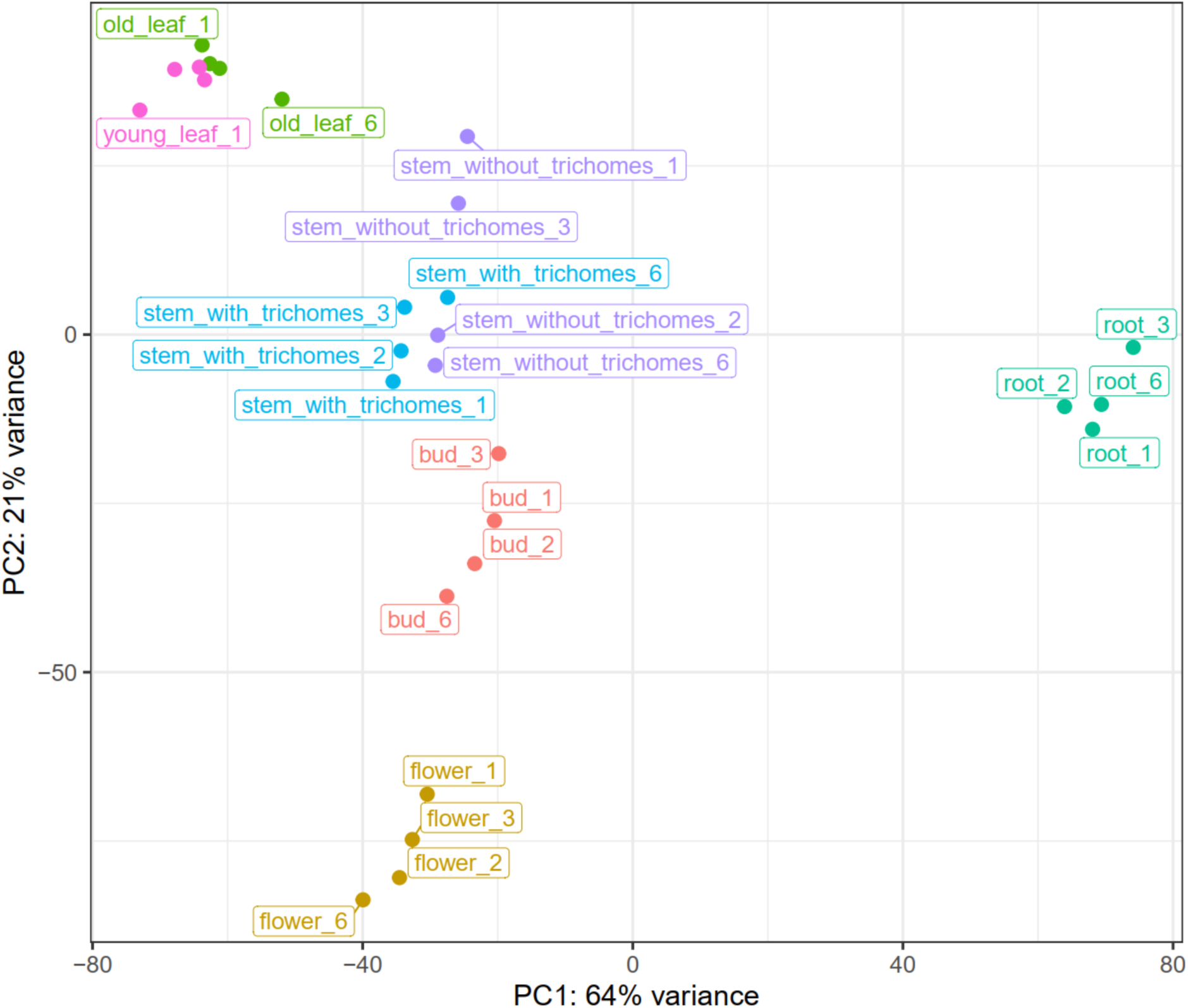
PCA analysis of the *S. nodosa* organ transcriptomes. The 4 replicates per organ used in the analysis are shown.

**Figure S2.**
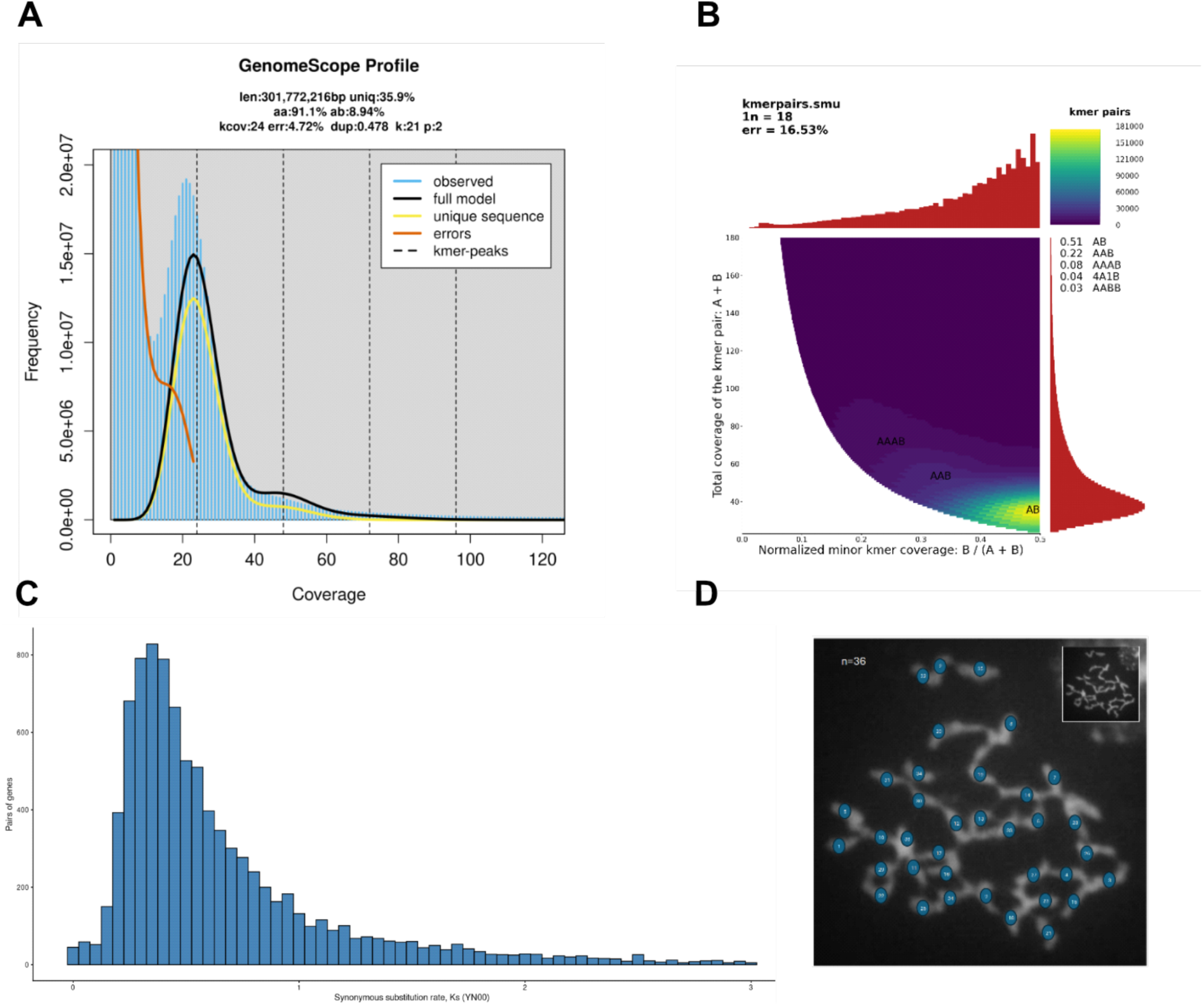
**A.** *S. nodosa* genome Genomescope profile. **B**. *S. nodosa* genome Smudgeplot analysis. **C**. Distribution of synonymous substitutions (Ks) of paralogous genes in the *S. nodosa* genome. **D**. Representative spreading and analysis of a metaphasic haploid cell from *S. nodosa* from unopened floral buds (chromosome count: n = 36). Young floral buds containing meiotic stages were collected and immediately fixed in freshly prepared ethanol:acetic acid (3:1, v/v). Buds were fully immersed in fixative and incubated overnight at 4 °C with gentle rotation. After fixation, samples were transferred to 70% ethanol for storage at 4 °C until use. Fixed buds were washed twice in water and twice in 10 mM citrate buffer (pH 4.5), followed by enzymatic digestion (0.3% cellulase, 0.3% pectolyase, and 0.3% cytohelicase) at 37 °C for 2 h to release pollen mother cells. Individual buds were then dissected on microscope slides and gently macerated to obtain a cell suspension. Chromosomes were spread by adding 60% acetic acid and incubating the suspension on a hot plate (45 °C) while stirring for 30-60 s to promote cell spreading. The preparation was then fixed by adding ethanol:acetic acid (3:1) around the droplet and allowed to air-dry. DNA was stained with DAPI (1 µg mL⁻¹) and meiotic chromosomes were visualized using fluorescence microscopy. Counting was done using ImageJ.

**Figure S3:**
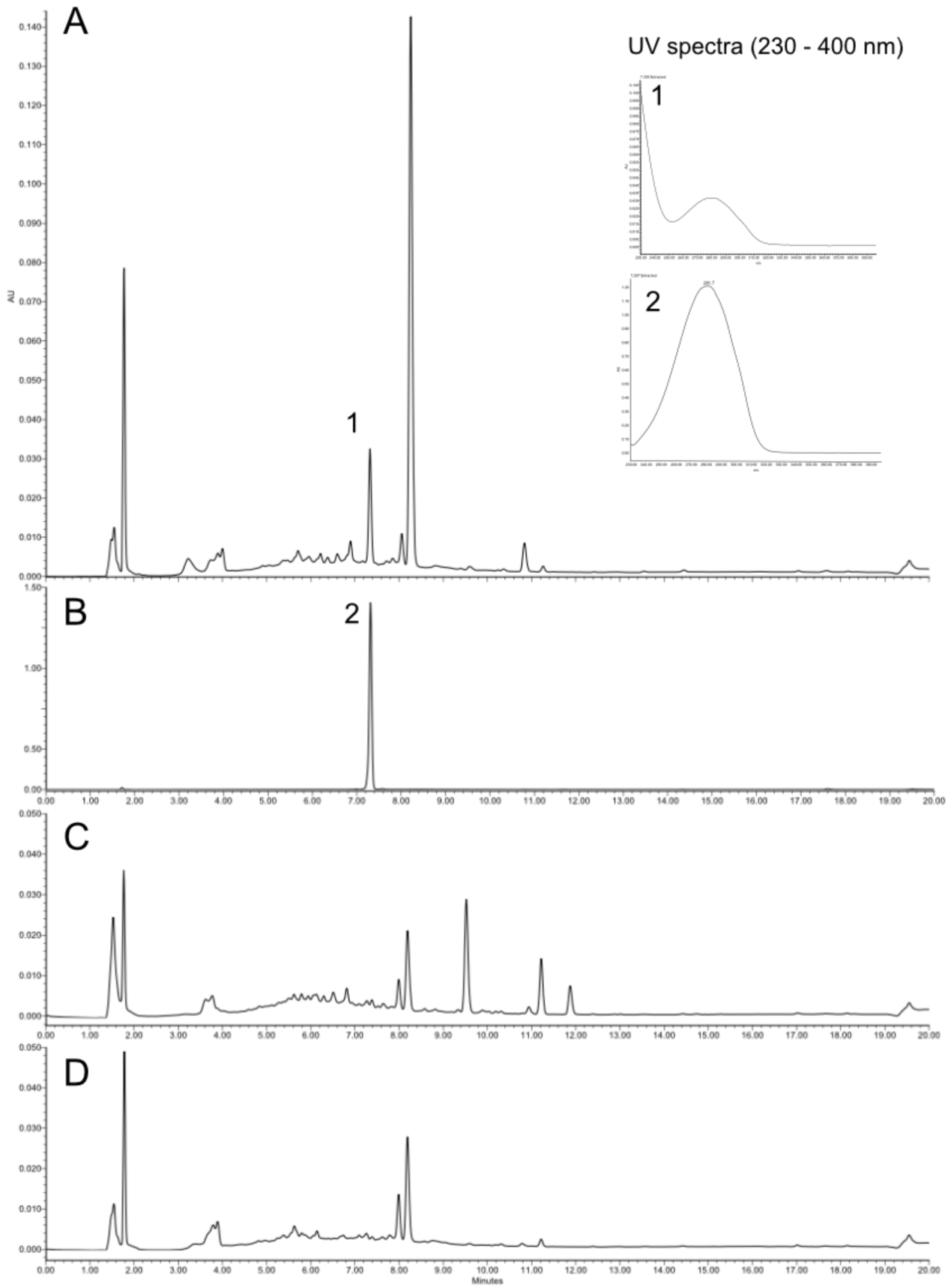
Semi-purified crude leaf protein extract activity test. A: Representative fraction assay in presence of harpagide and cinnamoyl-CoA. B: harpagoside standard. C. Assay with boiled protein fraction in presence of harpagide and cinnamoyl-CoA. D. Assay without cinnamoyl-CoA. Insert: UV spectrum of peak 1, reaction product, and 2, harpagoside standard, at RT = 7.3 min. All purification steps were carried at 4°C. 15 g of fresh *S. nodosa* leaves were ground in a blender with 100 mL pre-chilled protein extraction buffer A: TRIS pH = 8.0 50 mM, NaCl 300 mM, 10 mM β-mercaptoethanol (Sigma). After 30 min stirring, extracts were filtered on cloth to remove debris and centrifugated at 3500 g for 20 minutes. The recovered extract proteins were successively precipitated, first with addition of ammonium sulfate to 40% saturation, then after centrifugation and supernatant recovery, a second addition of ammonium sulfate to 80 % saturation was performed to precipitate the bulk of soluble proteins. After centrifugation, the recovered pellet was dissolved in 3 mL of buffer A. The resulting extract was then desalted on column to remove potential harpagoside carry-over, and activity was tested in the collected fractions. Activity was measured using 50 µL desalted extract, 5 µL TRIS 100 mM pH=8, 0.5 µL of 1 mM cinnamoyl-CoA, 1 µL of 30 mM harpagide. The reaction was performed for 30 minutes at 30° C and was stopped by addition of 100 µL 100% acetonitrile. Samples were centrifuged and transferred to vials for HPLC-UV analysis (see methods in main text).

**Figure S4:**
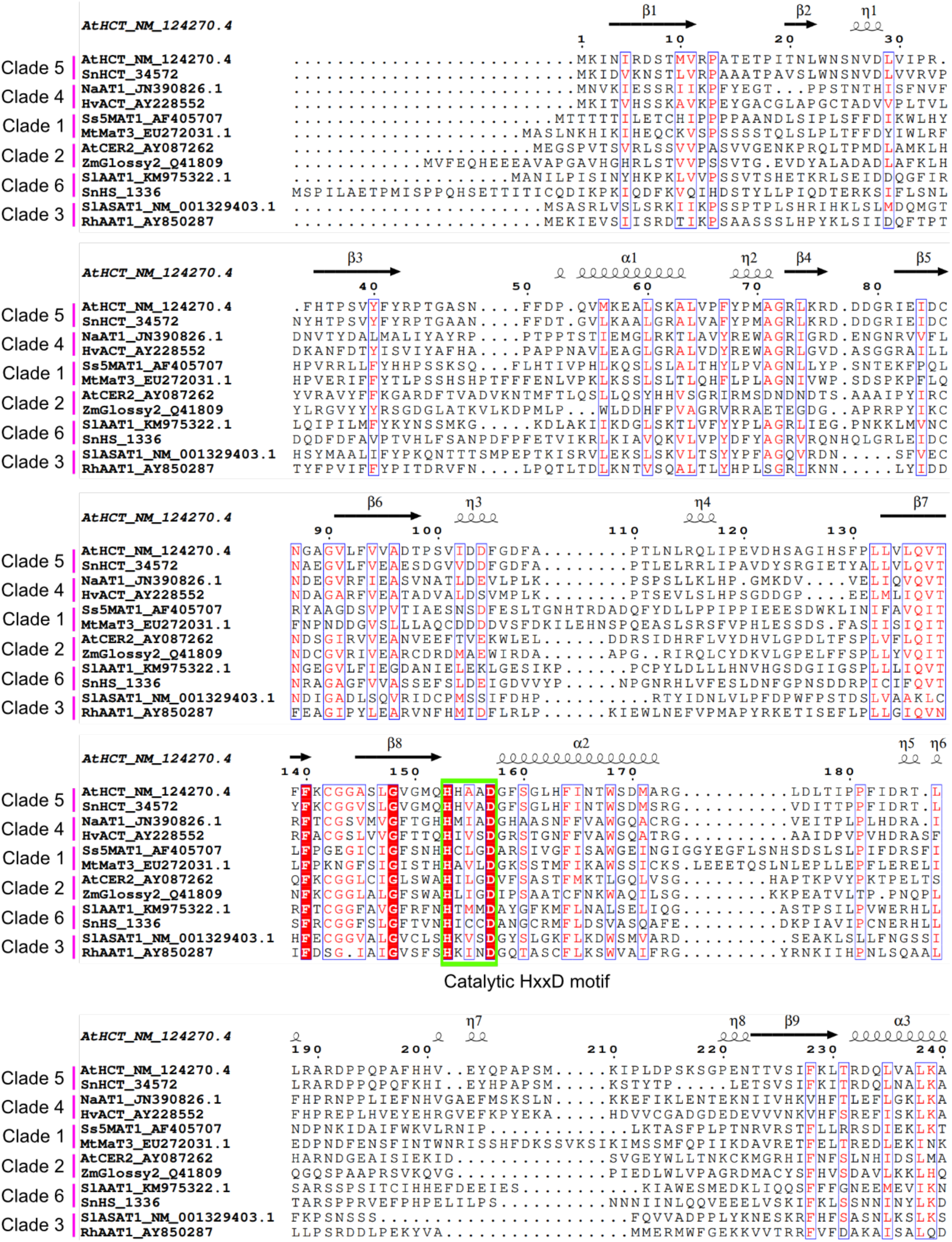
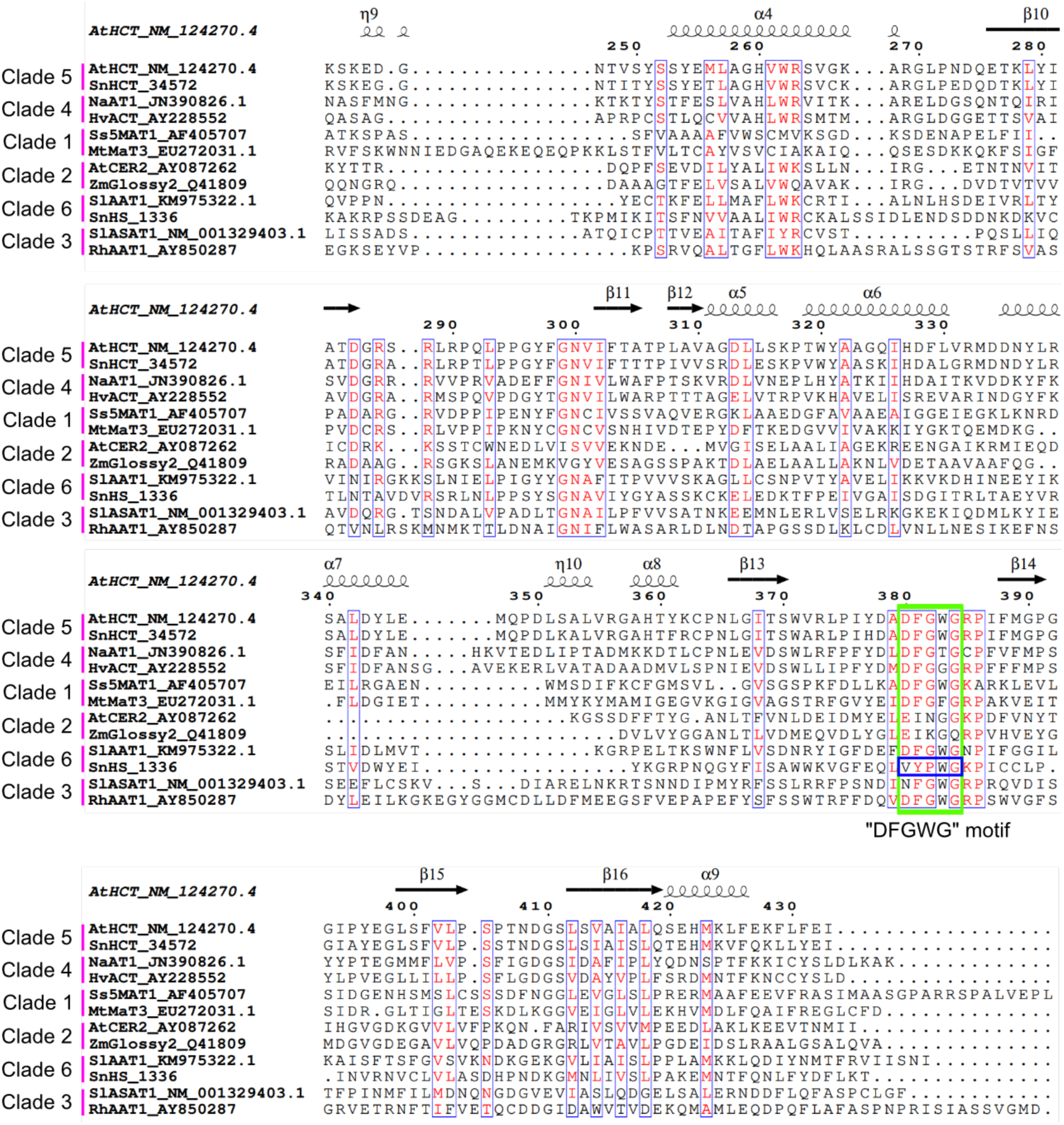
Amino acid sequences alignment of BAHDs representative of each clade (1 to 6, according to Kruse et al 2022). Catalytic site motif “HxxxD” and “DFGWG” motifs are shown in green boxes. The particular motif VYPWG from clade 6i branch 3 SnHS is shown in blue square. Note the typical DFGWG motif in the other representative clade 6 sequence shown (SlAAT1 from clade 6c).

**Figure S5.**
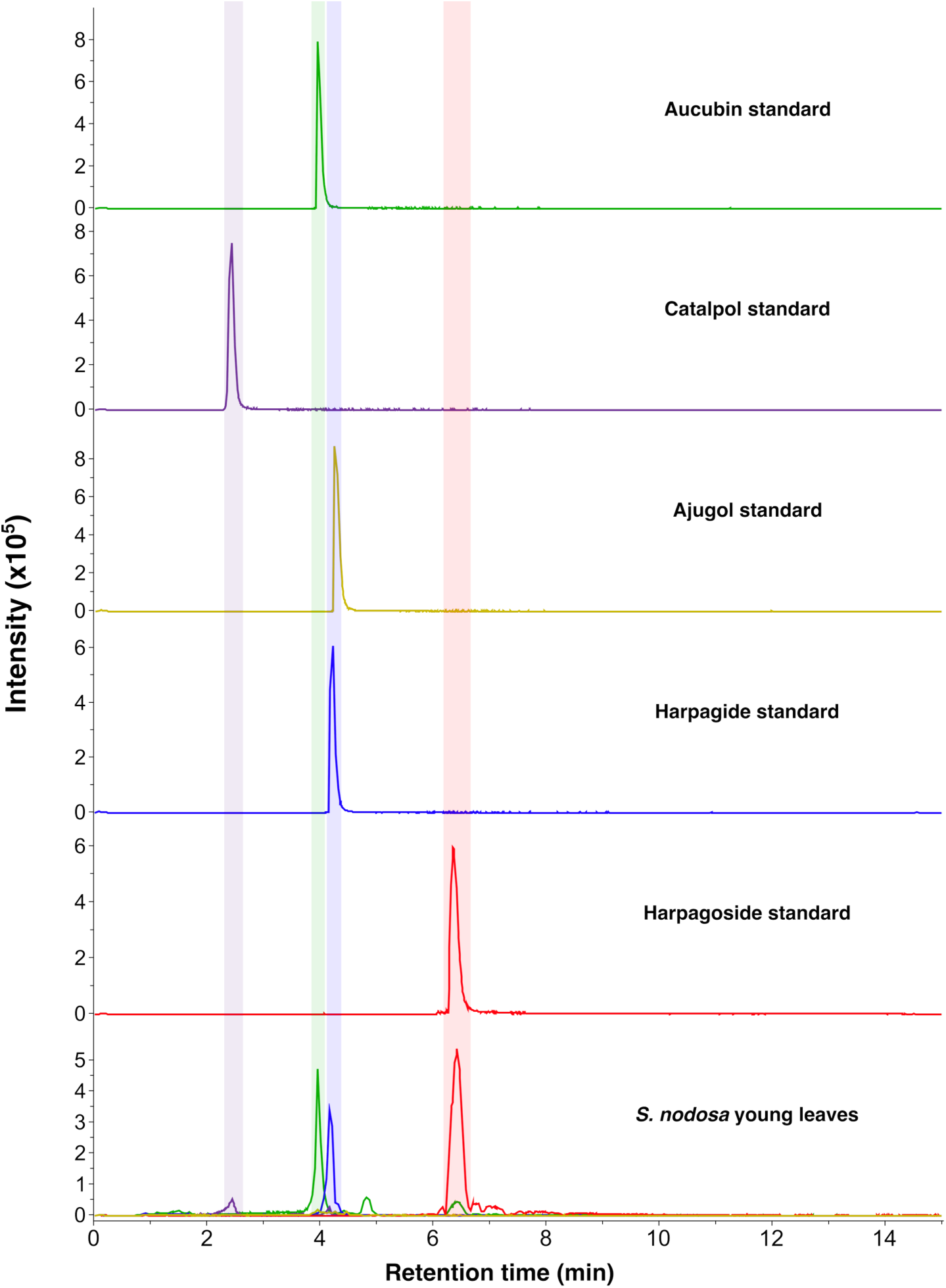
LC-MS/MS profiles of iridoid standards and *S. nodosa* young leaf extracts from a representative sample used for transcriptomics and metabolomics analysis. Intensity were recorded for aucubin (EIC: [M-H]^−^: 345.1191 ± 0.01; [M+HCOOH-H]^−^ = 391.1246 ± 0.01), catalpol (EIC: [M-H]^−^: 361.1140 ± 0.01; [M+HCOOH-H]^−^ = 407.1195 ± 0.01), ajugol (EIC: [M-H]^−^: 347.1348 ± 0.01; [M+HCOOH-H]^−^ = 393.1402 ± 0.01), harpagide (EIC: [M-H]^−^: 363.1297 ± 0.01; [M+HCOOH-H]^−^ = 409.1351 ± 0.01) and harpagoside (EIC: [M-H]^−^: 493.1715 ± 0.01; [M+HCOOH-H]^−^ = 539.1770 ± 0.01).

**Figure S6.**
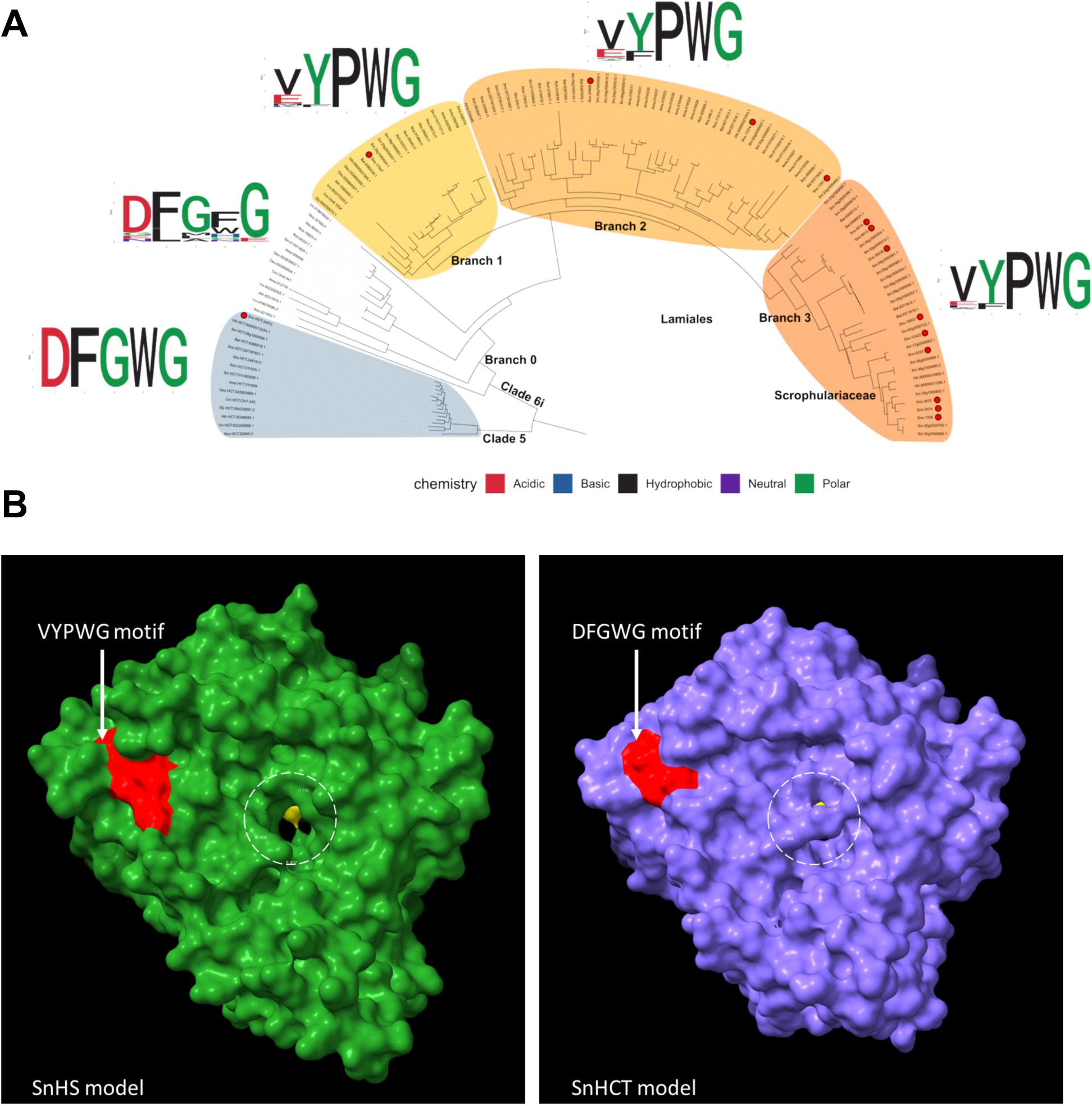
**A.** Phylogenetic tree of subclade 6i BAHD protein sequences and analysis of conserved protein motifs per branch. Clade 5 representative sequences were used to root the tree (blue). Red dots indicate *S. nodosa* sequences. Consensus (Weblogo) sequences of the canonical “DFGWG” motif are displayed; acidic amino acids are shown in red, basic in blue, hydrophobic in black, neutral in purple and polar in green. Two specific branches of Clade 6i enzymes, which produced multiple sequences in Lamiaceae (Branch 2) and Scrophulariaceae (Branch 3), show a highly conserved alternative sequence “VYPWG” for this motif. **B.** Models of *S. nodosa* SnHS and *S.nodosa* HCT homolog (SnHCT, clade 5) showing the subclade 6i specific VYPWG (in SnHS) and the typical DFGWG (in SnHCT) motifs in red. Conserved catalytic histidine residue (from HXXXD motif) is highlighted in yellow in the active site cavity (circled in white).

